# Constraints on the Efficiency of Electromicrobial Production

**DOI:** 10.1101/2020.06.23.167288

**Authors:** Farshid Salimijazi, Jaehwan Kim, Alexa Schmitz, Richard Grenville, Andrew Bocarsly, Buz Barstow

## Abstract

Electromicrobial production technologies (EMP) aim to combine renewable electricity and microbial metabolism. We have constructed molecular to reactor scale models of EMP systems using H_2_-oxidation and extracellular electron transfer (EET). We predict the electrical-to-biofuel conversion efficiency could rise to ≥ 52% with *in vivo* CO_2_-fixation. H_2_ and EET-mediated EMP both need reactors with high surface areas. H_2_-diffusion at ambient pressure requires areas 20 to 2,000 times that of the solar photovoltaic (PV) supplying the system. Agitation can reduce this to less than the PV area, and the power needed becomes negligible when storing ≥ 1.1 megawatts. EET-mediated systems can be built that are ≤ 10 times the PV area and have minimal resistive energy losses if a conductive extracellular matrix (ECM) with a resistivity and height seen in natural conductive biofilms is used. The system area can be reduced to less than the PV area if the ECM conductivity and height are increased to those of conductive artificial polymers. Schemes that use electrochemical CO_2_-fixation could achieve electrical-to-fuel efficiencies of almost 50% with no complications of O_2_-sensitivity.

## 1 Introduction

We are moving towards a world of plentiful renewable electricity [1–3]. However, to enable high penetration of renewables onto the grid, energy storage with a capacity thousands of times greater than today’s will be essential [4–7]. Despite significant advances in electrified transportation, the need for hydrocarbons in many applications like aviation could persist and even grow for decades to come [3]. Likewise, the need to sequester tens of gigatonnes of CO_2_ per year will also continue to grow [8, 9]. Electromicrobial production (EMP) technologies that combine biological and electronic components have the potential to use renewable electricity to power the capture and sequestration of atmospheric CO_2_ and convert it into high-density, non-volatile infrastructure-compatible transportation fuels [7, 10–12].

One of the most successful demonstrations of electromicrobial production to date, the Bionic Leaf [13, 14], is capable of converting solar power to the biofuel isopropanol at efficiencies exceeding the theoretical maximum of C_3_ and C_4_ photosynthesis [15, 16]. If coupled to some of the most efficient Si or GaAs solar photovoltaics (PVs) [17], the Bionic Leaf could even outperform cyanobacterial photosynthesis, the most efficient form found in nature [18]. However, the energy storage cost of photosynthesis is ultra-low [19, 20]. Any system that aims to supplant photosynthesis will need to dramatically exceed its efficiency, its convenience and preferably both.

To date, no one has systematically explored the constraints on the efficiency of electromicrobial production systems. Here we present a model for comparing the theoretical efficiencies of systems that supply electrons to metabolism by either H_2_-oxidation [13, 14, 21, 22] or through a conductive extracellular matrix (ECM) by extracellular electron transfer (EET) [23]; employ *in vivo* enzymatic, or *ex vivo* electrochemical CO_2_ fixation [24]; and transform fixed carbon to the biofuels isopropanol [25] or butanol [20–22]. This analysis lets us calculate the maximum theoretical efficiency of each system and gives a roadmap for how to achieve it.

## Theory, Results and Discussion

### General Theory

**Figs. 1A** and **1B** show simplified schematics of electromicrobial production systems with *in vivo* and *ex vivo* CO_2_-fixation, respectively. In **Fig. 1A** a microbe absorbs electricity to generate reducing equivalents needed to enzymatically fix CO_2_ *in vivo* and synthesize an energy storage molecule like polyhydroxybutyrate (PHB) or a hydrocarbon fuel. In **Fig. 1B** CO_2_ is first electrochemically reduced to a short-chain hydrocarbon like formate or formic acid *ex vivo* [27–29]. A microbe in the second cell absorbs electricity and further reduces and concatenates the initial fixation product to a longer-chain carbon compound. In both cases, electricity is absorbed into metabolism by either H_2_-oxidation (H_2_-mediated electromicrobial production; H_2_-EMP) or EET (EET-mediated electromicrobial production; EET-EMP) **Fig. 1C**. A complete list of symbols used in this article is included in **Table S1**.

We define the electrical energy conversion efficiency as the rate of energy storage molecule production, 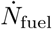, multiplied by the energy content per molecule, *E*_fuel_, relative to the total electrical power input,

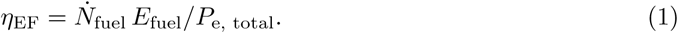

**Figure 1:**
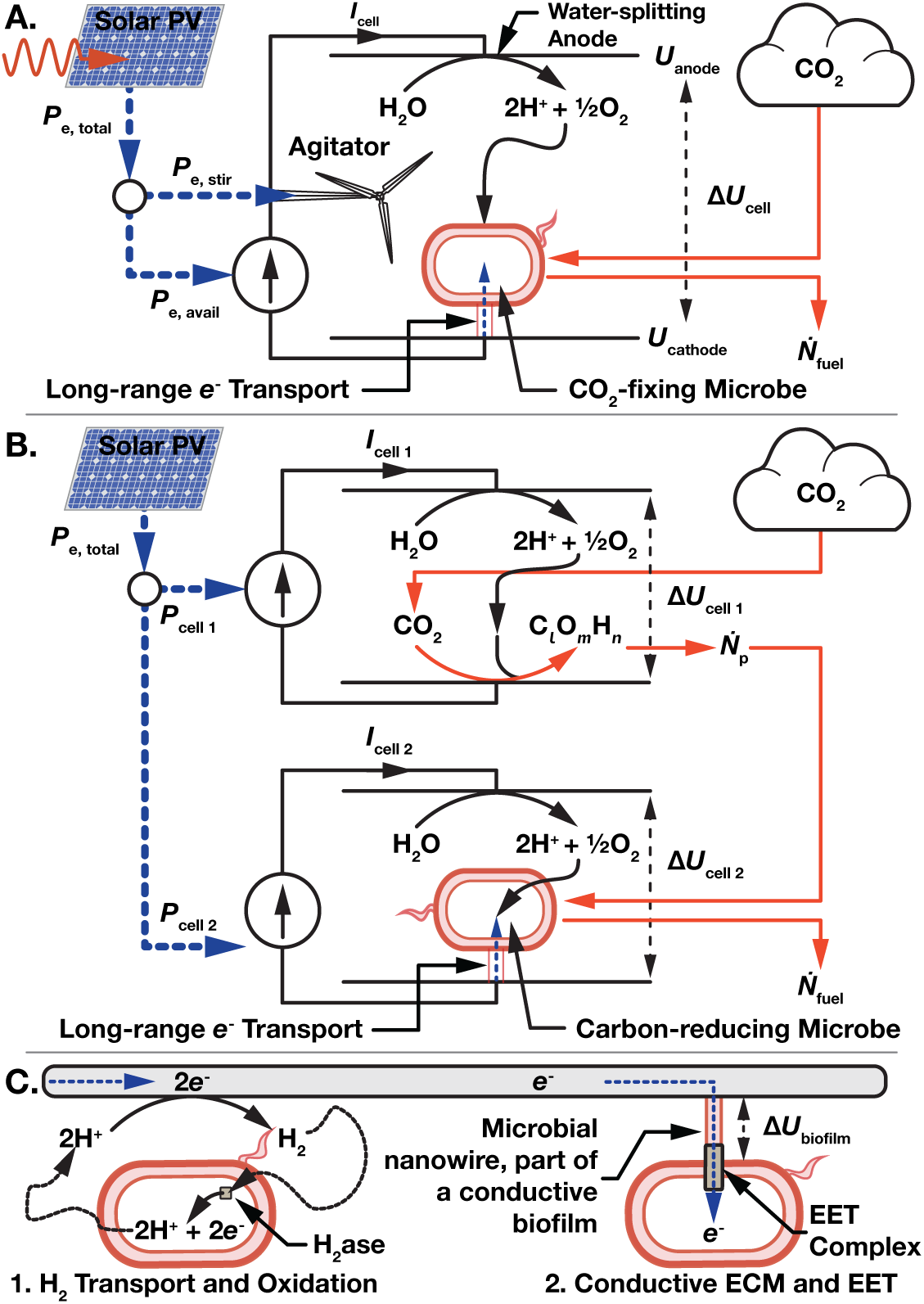
Overview of electromicrobial production technologies. (**A**) A microbe absorbs electrical power, *P*_e, avail_, through H_2_-oxidation or through a conductive extracellular matrix (ECM) by extracellular electron transfer (EET) to power CO_2_-fixation and biofuel production at a rate 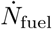. The total electrical power is used to drive a current, *I*_cell_, across a whole-cell voltage, Δ*U*_cell_, and can also be used to power an agitator. (**B**) The electrical power is split between two electrochemical cells. In the first CO_2_ is reduced to a short chain hydrocarbon like formic acid at a rate 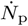. The primary fixation product is then concatenated in the second cell by a H_2_-oxidizing or electroactive microbe. (**C**) Electrons are transported to metabolism by either (1) diffusion or stirring of H_2_ and oxidation by a hydrogenase (H_2_ase) enzyme, or (2) across a conductive ECM and transport into an electroactive cell by a membrane-spanning EET complex. A bias voltage Δ*U*_biofilm_ is required to drive current across the ECM.

### H_2_-mediated Electromicrobial Production is Already Optimized but can be Improved by Swapping Out CO_2_-fixation

Estimating the efficiency of *in vivo* CO_2_-fixation (**Fig. 1A**) comes down to estimating 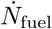 as a function of the electrical power available for electrochemistry, *P*_e, avail_; the voltage across the electrochemical cell, Δ*U*_cell_; and the number of electrons needed to generate the NAD(P)H, Ferredoxin (Fd), and ATP for synthesis of a single fuel molecule from CO_2_, *ν*_ef_ (*e* = elementary charge) (**SI Text 1**),

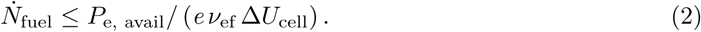

Therefore, the overall electrical to fuel efficiency for an *in vivo* CO_2_-fixation scheme,

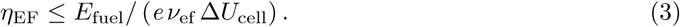

*ν*_ef_ can be estimated from molecular models of electron uptake. A schematic of the *Ralstonia eutropha* H_2_-oxidation machinery (used by references [13, 14, 21]) is shown in **Fig. 2A**. The low redox potential of H_2_ 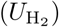 enables direct reduction of NADH by the cytosolic nickel-iron Soluble Hydrogenase (SH) (*R. eutropha* uses NADH rather than NADPH for CO_2_-fixation) [30, 31]. While the *R. eutropha* genome does not code for any Fd-reducing di-iron hydrogenases, these could be readily added to it [32–34]. Thus, the microbe simply has to oxidize a number of H_2_ molecules equal to the sum of NADH and Fd that it needs to synthesize a fuel molecule (the number of electrons needed is just double the number of H_2_).

**Figure 2:**
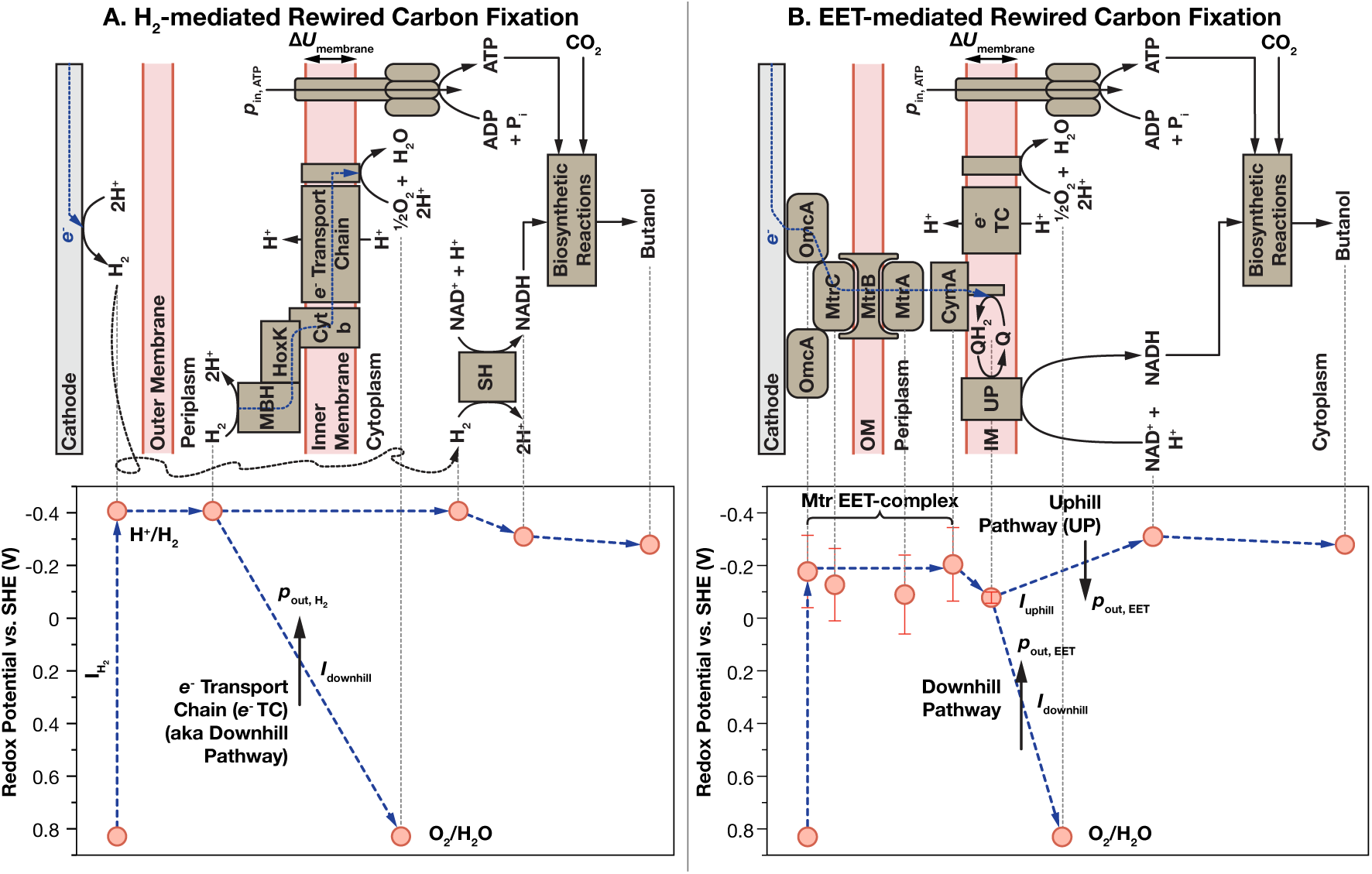
Energy landscapes for electromicrobial production. (**A**) In H_2_-mediated electromicrobial production, the incoming H_2_-current is used to directly reduce NAD(P)H or Ferredoxin (not shown), or is diverted into the conventional electron transport chain for ATP synthesis. For each electron sent downhill to reduce O_2_ and H^+^ to 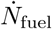 are pumped across the inner membrane. To synthesize an ATP molecule, *p*_in, ATP_ protons are released through the ATP synthase. (**B**) In Extracellular Electron Transfer (EET)-mediated electromicrobial production the incoming current is split between the uphill and downhill pathways. For each electron sent downhill, *p*_out, EET_ protons are pumped across the inner membrane. Midpoint redox potentials for the Mtr EET complex components are from Firer-Sherwood *et al.* [26].

ATP is generated by injection of electrons from H_2_-oxidation by the Membrane-Bound Hydrogenase (MBH) into the inner membrane electron transport chain [30, 31]; quantized energy transduction by proton pumping against the transmembrane voltage, Δ*U*_membrane_; reduction of a terminal electron acceptor at a redox potential *U*_Acceptor_; and further quantized energy transduction by proton release through the ATP synthase and ATP regeneration.

Therefore, the number of electrons needed to synthesize a single fuel molecule through H_2_-oxidation is (a full derivation is included in **SI Text 2**) (*ν*_f, NADH_, *ν*_f, Fd_, and *ν*_f, ATP_ are the number of NAD(P)H, Fd and ATP needed for synthesis of a single fuel molecule respectively),

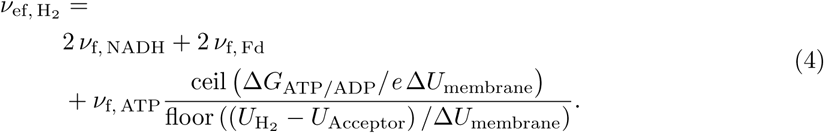

These equations are numerically solved with the REWIREDCARBON package using estimates for the NAD(P)H, ATP and Fd requirements for isopropanol and 1-butanol synthesis (**Fig. S2**) from CO_2_ fixed by the known natural CO_2_-fixation cycles and the synthetic CETCH cycle [35] in **Table S2**.

The biggest source of uncertainty in the efficiency estimate is the transmembrane voltage (Δ*U*_membrane_). At the time of writing we are unaware of any direct measurement of Δ*U*_membrane_ in *R. eutropha* or the electroactive microbe *Shewanella oneidensis*. Therefore, in **Fig. 3** we present a range of efficiency estimates for Δ*U*_membrane_ = 80 mV (BioNumber ID (BNID) 104082 [36]) to 270 mV (BNID 107135), with a central value of 140 mV (BNIDs 109774, 103386, 109775). Counterintuitively, the efficiency of H_2_-mediated electromicrobial production trends downwards, moving from plateau to plateau, with increasing transmembrane voltage. (**Fig. S1A**). While the amount of energy stored per proton is lower at lower Δ*U*_membrane_, energy quantization losses are also reduced.

**Figure 3:**
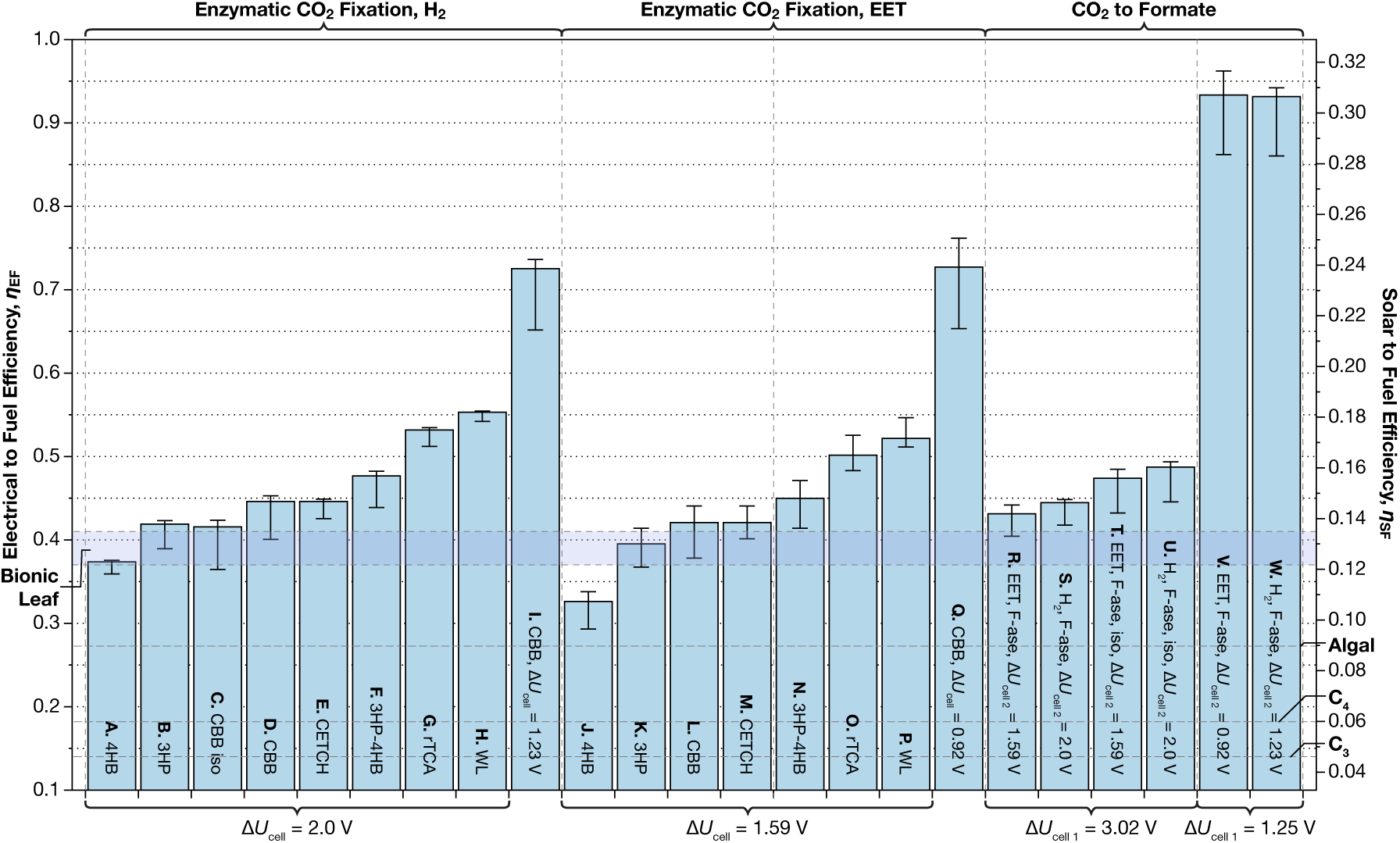
Projected lab-scale electrical and solar to biofuel efficiency of electromicrobial production schemes. The right axis is calculated by assuming a solar to electrical conversion efficiency of 32.9%, the maximum efficiency of a single junction Si solar PV [41]. Bars R to U assume a Faradaic efficency of CO_2_ to formate reduction of 80%, while bars V and W assume 100% Faradaic efficiency. Whole cell voltages were calculated from the minimum redox potentials of H_2_ and the Mtr EET complex [26] midpoint redox potentials, and from bias voltages reported by [13], [14], and [23]. Metabolic pathway data can be found in **Table S2**. All efficiences are for butanol production, except where noted as isopropanol (iso). This plot can be recreated with the `fig-co2fixation.py` program and `fig-co2fixation.csv` input file in the REWIREDCARBON package. 4HB = 4-hydroxybutyrate cycle; 3HP = 3-hydroxypropionate bicycle; CBB = Calvin-Benson-Bassham cycle; CETCH = (CoA)/ethylmalonyl-CoA/hydroxybutyryl-CoA; 3HP-4HB = 3-hydroxypropionate 4-hydroxybutyrate bicycle; rTCA = reductive tricarboxylic acid cycle; WL = Wood-Ljungdahl pathway.

This framework estimates the electron requirement for isopropanol and butanol synthesis by the Bionic Leaf (H_2_-EMP using the Calvin Cycle (CBB) for *in vivo* CO_2_-fixation) to be 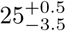 and 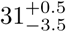 respectively. The maximum electricity to isopropanol conversion efficiency of the Bionic Leaf (Δ*U*_cell_ = 2 V [14]) is estimated to be 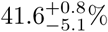 (**Bar C** in **Fig. 3**). This result just exceeds the maximum reported electrical to isopropanol efficiency of 39 *±* 2% [14]. This match suggests that CO_2_-fixation and biofuel synthesis in *R. eutropha* are already highly optimized.

How high could the efficiency go? Switching the product to butanol affords an improvement in H_2_-EMP efficiency to 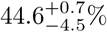 and a significant improvement in ease of product recovery (**Bar D** in **Fig. 3**). If the anode and cathode bias voltages could be reduced to zero, the efficiency of H_2_-EMP electrical to 1-butanol efficiency could rise as high as 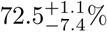 (**Bar I**). However, given the already low cobalt phosphate electrode overpotentials [37] in the Bionic Leaf, raising the efficiency by this route might be impractical.

Could the efficiency of EMP be increased by altering just the biological part of the system? Following intuition, electrical to fuel efficiency increases with decreasing NAD(P)H, ATP and Fd requirements for CO_2_ to biofuel conversion (**Fig. S3A-D**). The efficiencies of the six known naturally-occurring carbon fixation pathways and the synthetic CETCH pathway are shown in **Fig. 3**. The CETCH [35] cycle matches the efficiency of CBB (**Bar E**), while the naturally-occurring CO_2_-fixation cycles 3HP-4HB (**Bar F**), rTCA (**Bar G**) and WL (**Bar H**) all perform better than the Calvin cycle, raising the electrical to fuel efficiency as high as 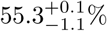.

While the rTCA cycle and Wood-Ljungdahl pathway are both typically found in anaerobic and micro-aerophilic organisms, recent advances in compartmentalization in synthetic biology [38–40] could enable the implementation of these highly efficient pathways in synthetic organisms that operate under ambient atmospheric conditions and enable use of O_2_ as a metabolic terminal electron acceptor.

### H_2_-mediated Electromicrobial Production Reaches Its Maximum Efficiency in Large Scale Systems

In principle, the efficiency of a electromicrobial production system could be independent of the specific activity of the carbon fixation pathway used (how many CO_2_ molecules are fixed each second by each gram of enzyme). Fixing more CO_2_ and storing more energy might simply require more cells operating in parallel. However, distributing electrical power through a H_2_ mediator could pose energetic, geometric and safety challenges [31]. To assess these challenges, we built models of H_2_-transport by diffusion and agitation.

The difficulty of H_2_-transport is determined by the number and volume of cells needed to store the H_2_-current, 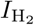, produced by the cell current (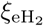 is the Faradaic efficiency of H_2_ production, typically close to 1),

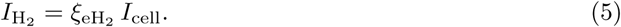

As hydrogenase enzymes are much faster than any carboxylating enzyme, the CO_2_ fixation rate is the limiting factor in electron demand per cell. The rate of electron uptake by each cell depends on the number of electrons, *ν*_ef_, and carbon atoms fixed, *ν*_Cf, fix_ (not just the number incorporated, *ν*_Cf_), to synthesize each fuel molecule; and the rate and number of carbon-fixing enzymes, *r*_fix_ and *ν*_fix_ (**SI Text 3**),

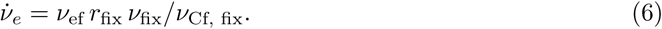

Thus, the total number and volume of cells needed to store the H_2_-current (n_cells_ is the cell density),

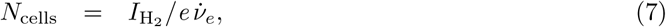

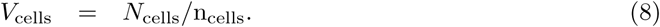

H_2_ could be transported by diffusion from the headspace of a reactor (where it is at a partial pressure 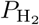) without any additional energy input into the system (**Fig. 4A**). In order to achieve the high concentration gradient needed to drive rapid diffusion of H_2_ (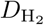 and 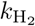 are the diffusion and solubility coefficients for H_2_ respectively), the cell culture has to be spread into a film with a height no greater than, and an area no less than (**SI Text 4**),

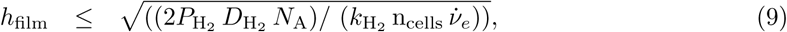

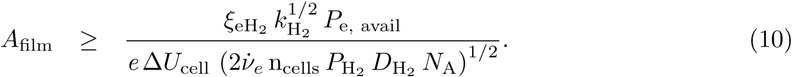

**Figure 4:**
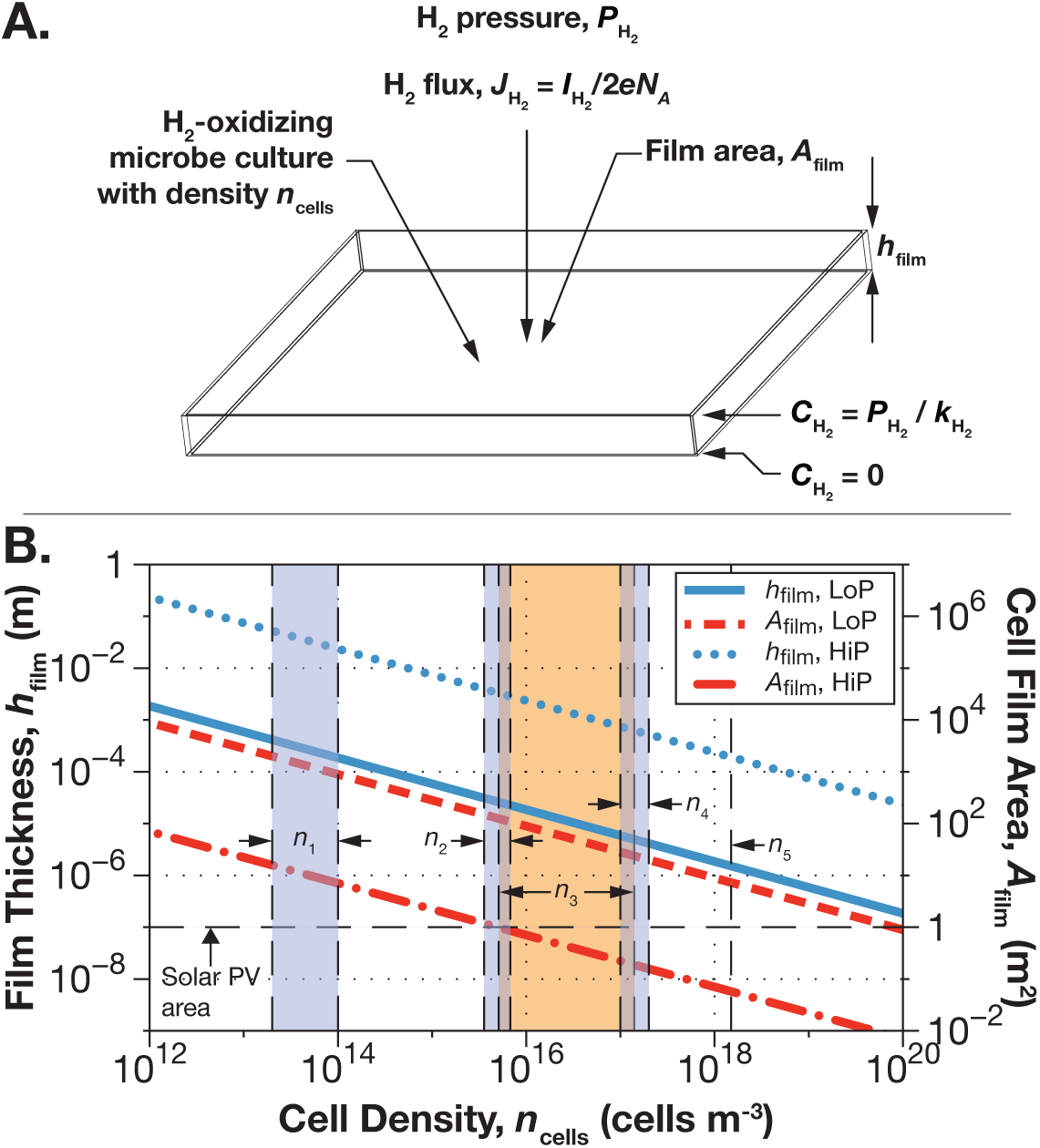
H_2_-transport by diffusion to enable scale up of H_2_-mediated electromicrobial production systems using the Calvin cycle (CBB) to convert CO_2_ to butanol. (**A**) Geometry for H_2_ mixing by diffusion. (**B**) Maximum height of cell culture that can be supplied with H_2_ by diffusion and corresponding area of culture needed to convert 330 W of electrical power (produced by a perfectly efficient 1 m^2^ single-junction Si solar PV illuminated by 1, 000× W of solar power) at H_2_ partial pressures of 5066 Pa (5% of atmospheric pressure; LoP) and 81 MPa (80% of 1000× atmospheric pressure; HiP). Five important cell density regimes are noted in panel **B**: *n*_1_: laboratory grown cultures of *E. coli* in exponential phase; *n*_2_: cyanobacteria grown to maximum density; *n*_3_: cultures of *E. coli* at saturating density; *n*_4_: H_2_-oxidizing microbes grown to maximum density; and *n*_5_: and saturating cultures of industrially-grown yeast (**SI Text 5**) and **Table S5**. Panel **B** can be recreated with the `fig-h2diffusion.py` programs and corresponding input file in the REWIREDCARBON package. To ease interpretation of panel **B** we have re-drawn this panel as two separate panels, each with a single curve representing the area and thickness of the cell culture film at each pressure in **Fig. S6**.

The area of a electromicrobial production system supplied by H_2_-diffusion scales linearly with input power while the film thickness remains the same. *R. eutropha* is typically grown under an atmosphere containing H_2_, O_2_ and CO_2_ at a ratio of 8:1:1 [42]. At the laboratory-scale, the H_2_ partial pressure is usually restricted to 5% of a total pressure of 1 atmosphere in order to reduce the risks of H_2_ explosion [42]. If supplied by a solar photovoltaic (PV), the area of the film relative to the solar PV area, *A*_PV_, will remain constant. A plot of film thickness and area versus cell culture density is shown for two systems supplied by a 1 m^2^ solar PV in **Fig. 4B**: the first with a headspace H_2_ partial pressure of 5,066 pascals (Pa) (5% of 1 atmosphere; O_2_ and CO_2_ will both be at a partial pressure of 633.25 Pa and the system will be balanced with N_2_), and the second with a H_2_ partial pressure of 81 × 10^6^Pa (80% of 1,000 atmospheres; O_2_ and CO_2_ will both be at a partial pressure of 10.1 × 10^6^Pa).

For the ambient pressure system, the film area (and potential footprint of the system) is greater than the area of the PV supplying it for even the highest cell densities seen in bio-industrial applications. At the highest reported autotrophic density for *R. eutropha* (density region **4**; *n*_4_ [43]), the film area is between 20 and 28 m^2^. The large film area requirement for H_2_-transport by diffusion at ambient pressures may not be insurmountable. Bioreactors with high internal areas but relatively small footprints could be constructed by stacking planar cell layers on top of one another, or using hollow fibers in which cells are immobilized on the walls of the fiber and reactant gases are flowed along its inner and outer surfaces [44].

Furthermore, by increasing the H_2_ partial pressure to 81 × 10^6^Pa, the cell film area can be reduced to 1 m^2^ by a density of ≈ 5 × 10^15^ cells m^−3^, inside the range of typical cyanobacterial cell densities (density region **2**; *n*_2_).

H_2_-diffusion systems could enable very high efficiency, but may come at the cost of high initial expenditure, complexity, maintenance, potential for H_2_ escape, and difficulty in removing product.

Intuitively, agitation allows H_2_-transport without the need for extreme system geometries, high pressures or both, at the expense of power input. The input power to the electrochemical cell is the total available electrical power, *P*_e, total_, minus any power needed to agitate the system,

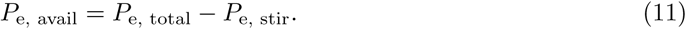

We considered a cylindrical stirred tank of cells that continuously distributes H_2_ supplied by a sub-surface pipe (**Fig. 5A**). We numerically solved a set of coupled equations linking H_2_ production, consumption, gas transfer rate, cell culture volume, and the power required for gas mixing through an iterative algorithm in the REWIREDCARBON package using a formalism compiled by Van’t Riet [45] until a self consistent set of solutions were found (**SI Text 6**). The solution to these equations for a system supplied with 330W of electrical power from a 1 m^2^ solar PV are plotted in **Figs. 5B** to **5D**.

**Figure 5:**
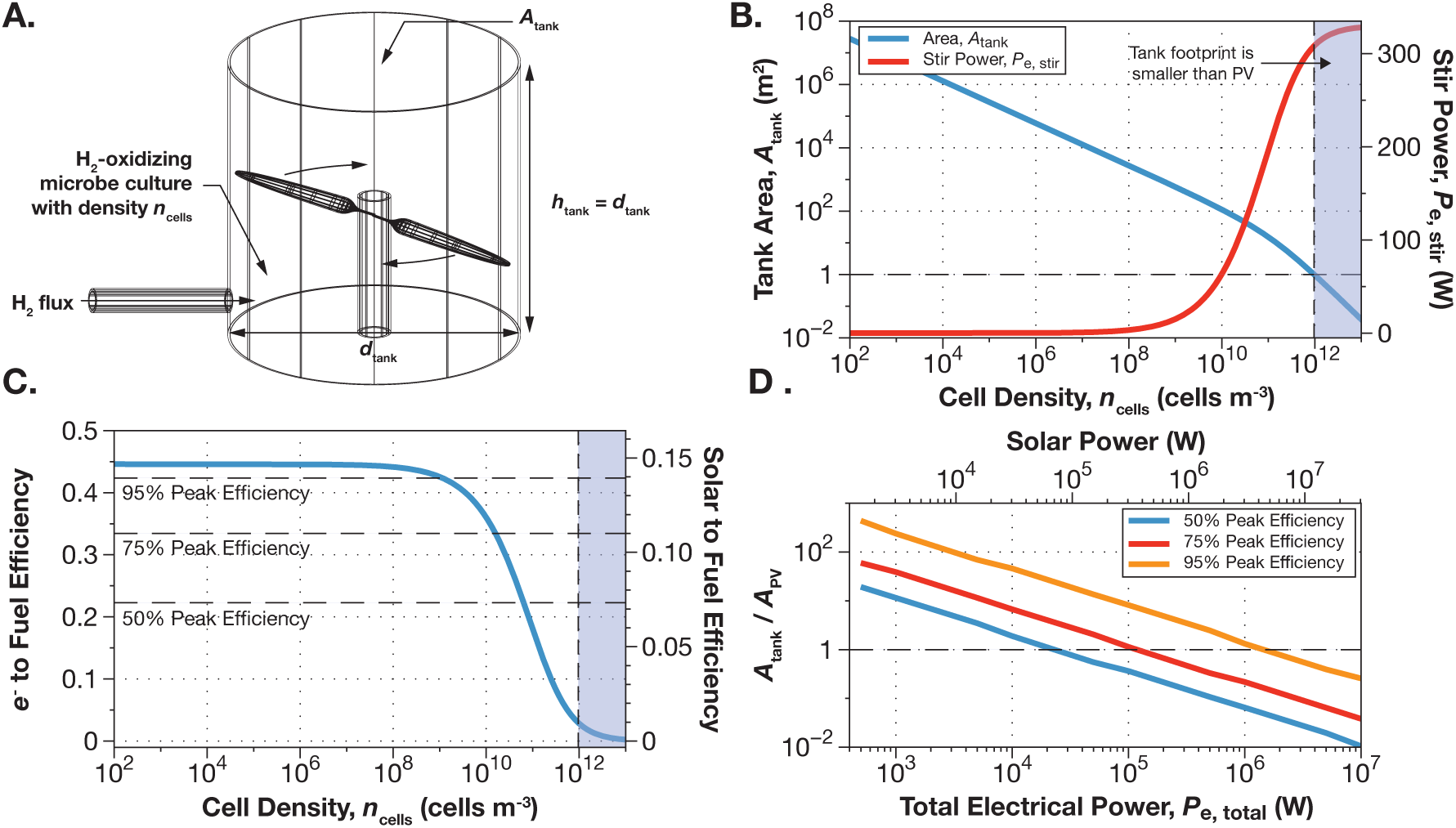
Scale up of H_2_-mediated electromicrobial production systems using the Calvin cycle (CBB) to convert CO_2_ to 1-butanol. (**A**) Geometry for mixing H_2_ by agitation. (**B**) As cell density is increased to reduce system footprint, the power required to mix H_2_ by agitation increases, eventually consuming all of the 330 W available to the system, (**C**) reducing the electricity to fuel efficiency to zero. (**D**) But, the system fooprint to PV area ratio at which the system achieves 50, 75 and 95% of its peak efficiency falls with increasing input power to the system (and solar PV area). Panels **B** to **D** in this plot can be recreated with the `fig-h2agitation-B` to `D.py` programs and the corresponding input files in the REWIREDCARBON package. Note that the cell densities shown here are much lower than those highlighted in **Fig. 4**.

At low cell densities and high system footprints (and hence volumes), the power required to transport H_2_ is low, while at low volumes the effort to stir is much greater (**Fig. 5B**). Intuitively, anyone who has grown cell culture understands that it is much easier to agitate a large cell culture (*e.g.* a 1 L flask) than a smaller culture (*e.g.* a 200 *μ*L well in a 384-well plate). This creates a conundrum, *P*_e, stir_ can be minimized, but at the expense of a tank footprint much larger *A*_PV_. Or, the tank footprint can be reduced to less than *A*_PV_, but at the expense of diverting more and more solar power to mixing H_2_ (**Fig. 5B**). This means that the efficiency of the electromicrobial production system (**Fig. 5C**) drops precipitously from its maximum potential value to almost zero as the footprint of the system is reduced to allow it to fit under the solar PV supplying it.

The footprint-efficiency dilemma can be resolved by operating at higher input power. We calculated the system footprint to PV area ratio (*A*_tank_*/A*_PV_) at which the system achieves 50%, 75%, and 95% of its maximum potential efficiency in **Fig. 5D**. For small scale systems (500 to 10^4^ W of solar power) footprints of 60× to 7× the area of the solar PV supplying them are required to achieve 75% of maximum efficiency. However, for large scales systems exceeding 1.1 × 10^5^ W of electrical power, the system footprint begins to shrink below that of the solar PV supplying it. Systems supplied by more than 1.1 × 10^6^ W of electrical power can achieve 95% of maximum efficiency and still have a footprint smaller than the solar PV supplying them.

### EET Matches the Efficiency of H_2_ and can Achieve High Efficiencies at Small Scales

Extracellular electron transfer (EET) could allow scale up of electromicrobial production through the use of a conductive biofilm to supply electrons to the cell (**Fig. 1C**). Electroactive microbes can transfer charge to, from and between external substrates like metals and even electrodes at distances up to a centimeter from the cell surface and use specialized metalloprotein complexes that connect the cell surface to the electron transport chain in the inner membrane (**Fig. 2B**) [46–49].

The energy landscape of EET has raised concerns about its use in electromicrobial production. The redox potentials of the membrane spanning cytochrome complex (Mtr in *S. oneidensis* at ≈ −0.1 V vs. the Standard Hydrogen Electrode (SHE) [50]) and the inner membrane electron carriers menaquinone (−0.0885 V [50]) and ubiquinone (0.1 V [50]) are too high to directly reduce NAD^+^ to NADH (−0.32 V [51]).

Nature suggests that the redox potential mismatch between the inner membrane and NAD^+^ is not insurmountable. Today, electroactive iron-oxidizing microbes are able to draw electrons from the oxidation of iron minerals at redox potentials from +0.7 to ≈ 0.1 V to power CO_2_-fixation and autotrophic metabolism [52, 53]. In the distant past it is thought that iron-oxidation powered the global carbon cycle [54]. It is speculated that an “uphill pathway” is able to lower the redox potential of electrons in the quinone pool to that of NAD^+^ [50].

Recently Rowe *et al.* [55] provided compelling evidence that a reverse electron transport chain providing an uphill pathway operates in S. oneidensis. While the the full complement of genes encoding this pathway remains unknown (although some parts have been found [55–58]), this pathway is proposed to operate by directing part of a cathodic current downhill in energy to a terminal electron acceptor and pumping protons across the inner membrane. The energy stored in the proton gradient is used to power NAD^+^ reduction and ATP production. A model for electron uptake by EET is shown in **Fig. 2B**.

Due to the need to sacrifice some current to generate a proton gradient for NAD^+^ (and possibly Fd) reduction, the number of electrons needed to produce the NADH, Fd and ATP for synthesis of a single fuel molecule through EET is higher than in H_2_-oxidation (a full derivation is included in **SI Text 7**),

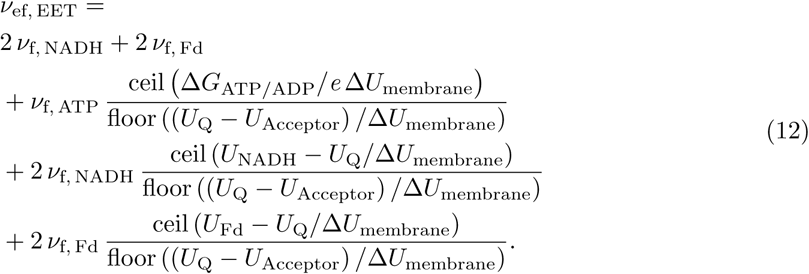

However, counterintuitively, EET-mediated electromicrobial production is not dramatically less efficient than H_2_-mediated electromicrobial production (**Fig. 3**). While the number of electrons needed to produce a molecule of fuel is higher, the whole-cell voltage in an EET-mediated system is lower than in a H_2_-mediated system (Δ*U*_cell_ ≥ 1.23 V for H_2_ but only ≥ 0.92 V for EET) as the redox potential of Mtr is much lower than H_2_ [26]. Furthermore, the bias voltages at lab-scale remain approximately the same [7], meaning more total current is available to an EET-mediated system. However, EET-mediated electromicrobial production is approximately twice as sensitive to changes in transmembrane voltage than a H_2_-mediated system (**Fig. S1**).

The scale up of EET-mediated electromicrobial production is potentially much easier than H_2_-EMP. We built a model of scale up for an EET-mediated system assuming that the dominant source of overpotential is the resistivity of the biofilm. We assumed that the biofilm could be modeled as an Ohmic resistor, so that the bias voltage needed to transport electrons across it is,

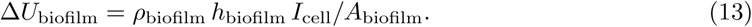

We developed a set of five coupled equations to solve for the cell current *I*_cell_, the bias voltage needed to drive current across the biofilm Δ*U*_biofilm_, the area of the biofilm *A*_biofilm_, the total number of cells in the biofilm *N*_cells_, and the volume of the biofilm *V*_biofilm_ in **SI Text 8**. These equations were solved numerically and the results shown in **Fig. 6**. Unlike agitation based systems, the energy cost of electron transport by EET scales linearly with system size: for a given biofilm resistivity, the ratio of the areas of the biofilm and the solar panel supplying it with electricity remains constant. Moreover, there is no obvious penalty for operating small-scale systems as there is with agitation.

**Figure 6:**
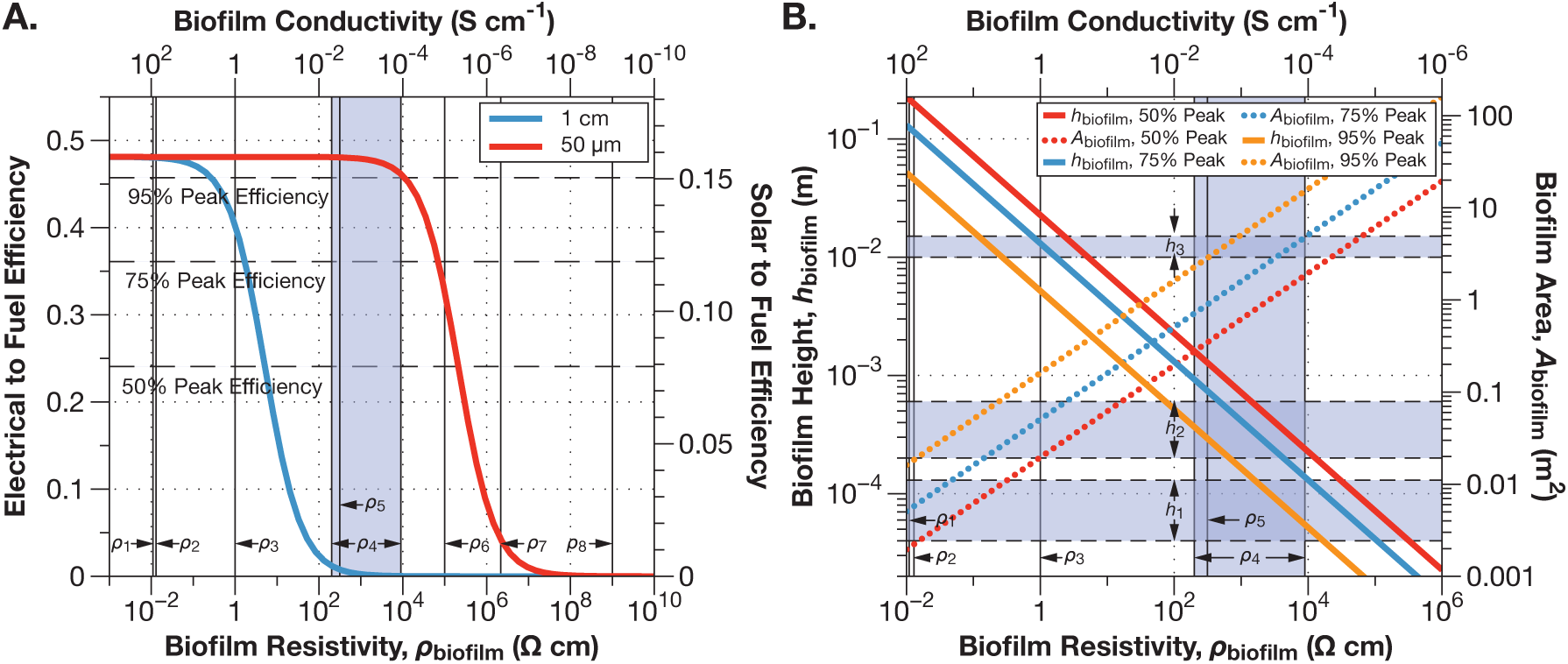
Biofilm resistivity determines efficiency losses in the scale-up of EET-mediated electromicrobial production. The system shown here has an anode bias voltage of 0.47 V, fixes CO_2_ with the Calvin cycle and produces butanol (**A**) The electrical to fuel efficiency of a electromicrobial production system drops after a threshold resistivity is reached. The thicker the biofilm, the earlier this drop occurs. (**B**) Maximum biofilm thickness and minimum area needed to achieve 50, 75 and 95% of peak efficiency. This plot can be recreated with the `fig-EETscaleup-A.py` and `B.py` programs and corresponding input files in the REWIREDCARBON package. Representative conductive matrix resistivities and heights: *ρ*_1_: high conductivity polypyrrole; *ρ*_2_: individual cable bacteria filaments; *ρ*_3_: individual *S. oneidensis* nanowires; *ρ*_4_: bulk *G. sulfurreducens* and *S. oneidensis* biofilm resistivities; *ρ*_5_: polypyrrole conductive matrix for *S. oneidensis*; *ρ*_6_: bulk *E. coli* biofilm; *ρ*_7_: HBr doped polyaniline; *ρ*_8_: low conductivity polypyrrole; *h*_1_: *G. sulfurreducens* biofilms; *h*_2_: polypyr-role conductive matrix for *S. oneidensis*; *h*_3_: cable bacteria biofilms and individual filaments (**SI Text 9**) and **Tables S3** and **S4**). To ease interpretation of panel **B** we have re-drawn this panel as three separate panels, each with a single curve representing the area and thickness of the biofilm at each efficiency in **Fig. S7**.

At low resistivities (high conductivities) the biofilm overpotential is small, allowing a conductive matrix system to achieve close to its maximum possible efficiency, set only by the thermodynamic minimum voltages and any non-biofilm bias in the system (**Fig. 6A**). However, above a critical resistivity, the efficiency drops precipitously. For a 50 *μ*m thick film, the efficiency starts to drop below 95% of maximum at a resistivity of ≈ 10^5^ Ω cm, considerably higher than the commonly reported resistivities of *Geobacter sulfurreducens* and *S. oneidensis* biofilms (*ρ*_4_ in **Fig. 6A, SI Text 9**) [59–61]. Note that the peak efficiency shown in **Fig. 6A** exceeds that shown in **Fig. 3** Bar L as we assume only anode bias.

As the resistivity of the conductive matrix increases, its thickness must decrease and its area increase in order to maintain a given efficiency. In contrast to a 50 *μ*m film, a 1 cm thick film suffers a drop in efficiency to 50% of maximum at a resistivity of only ≈10 Ω cm, well below the resistivity range of *G. sulfurreducens* and *S. oneidensis* biofilms [59–61], but above the reported resistivities of individual *S. oneidensis* nanowires (*ρ*_3_ in **Fig. 6A**) [62] and individual filaments produced by the cable bacterium *Thiofilum facile* (*ρ*_2_ in **Fig. 6A**) [63].

**Fig. 6B** shows the maximum conductive matrix thickness and minimum area able to achieve a given fraction of peak efficiency as a function of resistivity. If 50% of peak efficiency is acceptable, then the biofilm area can be constrained to 1 m^2^ (equal to that of the solar PV supplying it) if the biofilm resistivity is 2, 650 Ω cm, well within the range of *G. sulfurreducens* and *S. oneidensis* biofilm resistivities. However, the corresponding film thickness is 440 *μ*m, about 3× the height of most commonly observed *G. sulfurreducens* and *S. oneidensis* biofilms (although Renslow *et al.* did observe *S. oneidensis* films as thick as 450 *μ*m). However, artificial polypyrrole conductive ECMs have been produced that are as thick as 600 *μ*m, and have resistivities as low as 312 Ω cm (*ρ*_5_ in **Fig. 6**). Were the film area increased to 3.4 m^2^, the film thickness could be reduced to 130 *μ*m, within the range of commonly observed *G. sulfurreducens* and *S. oneidensis* biofilm thicknesses. The biofilm resistivity would only need to be 29, 000 Ω cm, above that of many conductive biofilms, perhaps allowing some conductivity to be sacrificed to enable increased CO_2_ inflow or biofuel outflow.

On the other hand, if a thickness of 130 *μ*m and resistivity of 1, 600 Ω cm are simultaneously achievable, 95% of peak efficiency can be achieved if a 6.4 m^2^ biofilm area is acceptable. If a 1 m^2^ biofilm with a resistivity 38 Ω cm and a thickness of 830 *μ*m could be produced, 95% of peak efficiency could be achieved.

If a biofilm could be produced with a 1 cm thickness (within the range of biofilm thickness produced by cable bacteria; *h*_3_ in **Fig. 6**), a resistivity of 5 Ω cm (above the resistivity of individual *S. oneidensis* nanowires, and well above that of individual *T. facile* filaments, but below that of the minimum resistivity calculated by Polizzi *et al.* of 30 Ω cm [64]), and an area of only 0.044 m^2^ then 50% of maximum efficiency could be achieved. If a biofilm of 1 cm thickness, with a resistivity of 0.26 Ω cm, and an area of 0.079 m^2^, 95% of peak efficiency could be achieved.

Finally, if 95% of peak efficiency were desired, but only a thin biofilm of 55 *μ*m with a high resistivity of 8, 952 Ω cm could be produced, then an area of 15 m^2^ would be required.

### Electrochemical CO_2_ Fixation Could Allow Very High Electricity to Fuel Conversion Efficiencies

H_2_-oxidation and EET could be an important complement to electrochemical CO_2_-fixation technologies. Current electrochemical CO_2_-fixation systems typically produce compounds with no more than two carbons that are often not completely reduced [27]. By contrast, most drop-in fuels require at least 2 to 3 carbons, with 8 electrons each.

Li *et al.* demonstrated the reduction of formate to isobutanol and 3-methyl-1-butanol (3MB) by the H_2_-oxidizing microbe *R. eutropha* [21]. While this work relied upon oxidation of formate to CO_2_ and subsequent re-fixation by RuBisCO, recent advances in artificial computational metabolic pathway could enable enzymatic transformation without reliance upon this bottleneck [65, 66].

The efficiency of electrochemical CO_2_-fixation electromicrobial production schemes is set by the number of electrons *ν*_e, add_ needed to produce the NAD(P)H, Fd and ATP needed to transform the primary fixation product to a biofuel; the charge needed to synthesize the primary electro-chemical CO_2_-fixation product, *e ν*_ep_; the number of carbons in each primary fixation product, *ν*_Cp_; the Faradaic efficiency of the first electrochemical reaction, *ξ*_I1_, (while we are calculating an upper limit on efficiency we have rarely seen *ξ*_I1_ > 0.8 [27]); the efficiency of carbon transfer to the second cell *ξ*_C_; and the Faradaic efficiency in the second cell *ξ*_I2_ (**SI Text 10**),

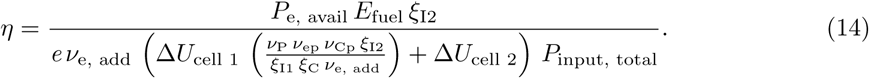

Even with only 80% Faradaic efficiency for the conversion of CO_2_ to formate, the electrical energy to butanol conversion efficiency of the formolase artificial metabolic pathway [65] powered by either H_2_-oxidation or EET exceeds all fully enzymatic CO_2_-fixation pathways with the exception of the rTCA cycle and Wood-Ljungdahl pathway **Fig. 3**, and suffers no complications of O_2_-sensitivity.

## Conclusions

What combination of electron uptake, electron transport, and carbon fixation is the best for electromicrobial production? The model of electromicrobial production lets us sketch out a roadmap for how to proceed with the technology. We outline 10 possible development and deployment scenarios that could be pursued in the near and further future in **Table 1** along with their advantages, disadvantages, and suggested niche.

**Table 1:**
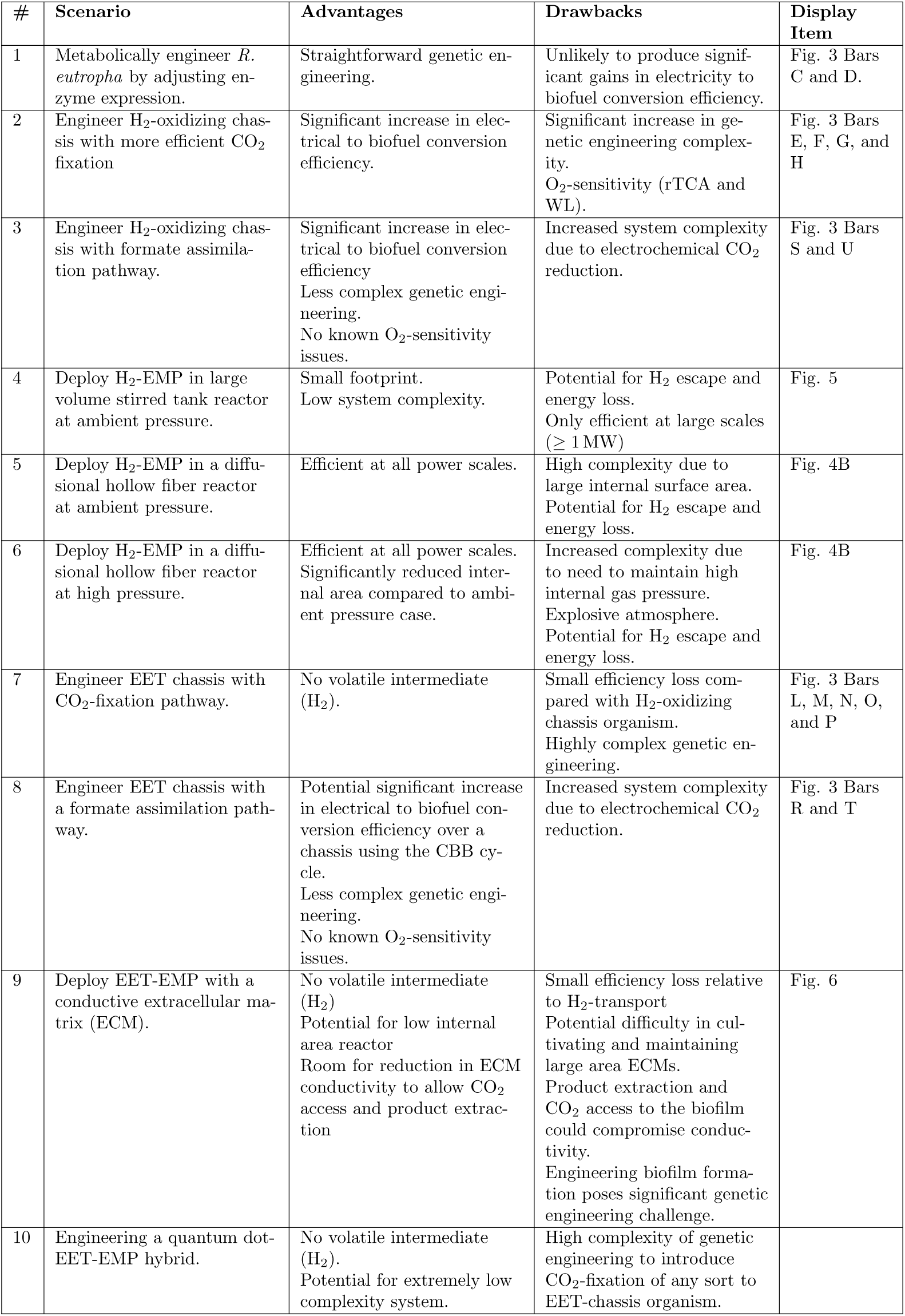
Future research and development, and deployment scenarios for electromicrobial production. ECM = Extra-Cellular Matrix.

| # | Scenario | Advantages | Drawbacks | Display Item |
| --- | --- | --- | --- | --- |
| 1 | Metabolically engineer *R. eutropha* by adjusting enzyme expression. | Straightforward genetic engineering. | Unlikely to produce significant gains in electricity to biofuel conversion efficiency. | Fig. 3 Bars C and D. |
| 2 | Engineer H<sub>2</sub>-oxidizing chassis with more efficient CO<sub>2</sub> fixation | Significant increase in electrical to biofuel conversion efficiency. | Significant increase in genetic engineering complexity. O<sub>2</sub>-sensitivity (rTCA and WL). | Fig. 3 Bars E, F, G, and H |
| 3 | Engineer H<sub>2</sub>-oxidizing chassis with formate assimilation pathway. | Significant increase in electrical to biofuel conversion efficiency<br>Less complex genetic engineering.<br>No known O<sub>2</sub>-sensitivity issues. | Increased system complexity due to electrochemical CO<sub>2</sub> reduction. | Fig. 3 Bars S and U |
| 4 | Deploy H<sub>2</sub>-EMP in large volume stirred tank reactor at ambient pressure. | Small footprint.<br>Low system complexity. | Potential for H<sub>2</sub> escape and energy loss.<br>Only efficient at large scales ($\geq 1$ MW) | Fig. 5 |
| 5 | Deploy H<sub>2</sub>-EMP in a diffusional hollow fiber reactor at ambient pressure. | Efficient at all power scales. | High complexity due to large internal surface area.<br>Potential for H<sub>2</sub> escape and energy loss. | Fig. 4B |
| 6 | Deploy H<sub>2</sub>-EMP in a diffusional hollow fiber reactor at high pressure. | Efficient at all power scales.<br>Significantly reduced internal area compared to ambient pressure case. | Increased complexity due to need to maintain high internal gas pressure.<br>Explosive atmosphere.<br>Potential for H<sub>2</sub> escape and energy loss. | Fig. 4B |
| 7 | Engineer EET chassis with CO<sub>2</sub>-fixation pathway. | No volatile intermediate (H<sub>2</sub>). | Small efficiency loss compared with H<sub>2</sub>-oxidizing chassis organism.<br>Highly complex genetic engineering. | Fig. 3 Bars L, M, N, O, and P |
| 8 | Engineer EET chassis with a formate assimilation pathway. | Potential significant increase in electrical to biofuel conversion efficiency over a chassis using the CBB cycle.<br>Less complex genetic engineering.<br>No known O<sub>2</sub>-sensitivity issues. | Increased system complexity due to electrochemical CO<sub>2</sub> reduction. | Fig. 3 Bars R and T |
| 9 | Deploy EET-EMP with a conductive extracellular matrix (ECM). | No volatile intermediate (H<sub>2</sub>)<br>Potential for low internal area reactor<br>Room for reduction in ECM conductivity to allow CO<sub>2</sub> access and product extraction | Small efficiency loss relative to H<sub>2</sub>-transport<br>Potential difficulty in cultivating and maintaining large area ECMs.<br>Product extraction and CO<sub>2</sub> access to the biofilm could compromise conductivity.<br>Engineering biofilm formation poses significant genetic engineering challenge. | Fig. 6 |
| 10 | Engineering a quantum dot-EET-EMP hybrid. | No volatile intermediate (H<sub>2</sub>).<br>Potential for extremely low complexity system. | High complexity of genetic engineering to introduce CO<sub>2</sub>-fixation of any sort to EET-chassis organism. |  |

This work shows that H_2_-EMP using the Calvin cycle [13, 14], is already highly optimized. This means that engineering the host microbe (*e.g. R. eutropha*) by adjusting expression levels of enzymes already encoded in the genome or changing the transmembrane voltage are unlikely to produce gains of more than a few percentage points in electricity to biofuel conversion efficiency.

One genetic engineering route to increased electrical to biofuel conversion efficiency (from ≈ 40% to as high as 55% at lab scales) is replacement of the familiar Calvin cycle with any one of the CETCH, 3HP-4HB, rTCA or WL CO_2_-fixation pathways. This is approach is not for the faint hearted. However, recent impressive progress in engineered the Calvin cycle into *E. coli* makes this a tantalizing possibility [67, 68]. Furthermore, the need to use O_2_ as a terminal electron acceptor to achieve maximum efficiency means that the O_2_-sensitivity of the rTCA and WL pathways will need to be mitigated by developing O_2_-tolerant versions of currently O_2_-sensitive enzymes in these pathways, or sequestering these enzymes inside O_2_-impermeable compartments inside the cell.

An alternative route to significantly enhanced efficiency is dispense with *in vivo* CO_2_-fixation and replace it with *ex vivo* electrochemical CO_2_ reduction and *in vivo* formate assimilation. This approach is much more genetically tractable and achieves efficiency gains comparable to replacing the Calvin cycle wih the rTCA cycle. Additionally, there is room for further improvement as new artificial pathways for processing electrochemically fixed CO_2_ are invented. However, this approach adds further system complexity and potential cost.

The optimization of H_2_-EMP with the Calvin cycle raises the question: is it time to take it out of the lab? Agitation is the most mature, lowest cost, and most easily implemented technology for electron transport considered in this article. However, the high energy cost of stirring small volumes means that the smallest increment of storage that can be built is ≈ 1 MW, about the size of a large solar farm. This is very large relative to residential storage needs (the average American home uses electrical energy at the rate of about 1.3 kW), but tiny compared to the production needs for aviation fuel (when converted to jet fuel with ≈ 50% efficiency 1 MW corresponds to ≈ 50 L hr^−1^. A 787-9 consumes fuel at the rate of ≈ 7, 000 Lhr^−1^).

Its not clear that H_2_-EMP will ever take on batteries for home energy storage. H_2_-EMP could operate very efficiently at a small power scale if H_2_ is transported by diffusion. However, this approach demands a high internal area reactor. This problem can be ameliorated by operating at high H_2_ pressure, but it is likely that this will increase cost, and incur significant safety risks. We would be foolish if we dismissed this approach outright, but we believe this analysis highlights significant technology risks.

Counter to intuition, the efficiency of EET-EMP using a reverse electron transport chain could almost match that H_2_-mediated electromicrobial production with laboratory overpotentials. Additionally its possible to grow conductive ECMs with sufficiently high conductivities and thicknesses that a high-efficiency, low-footprint, low internal area system could be produced with the microbes we already have available today. In principle, EET-EMP coupled to a self-assembled conductive extracellular matrix (ECM) could reduce construction costs; allow us to dispense with volatile intermediates like H_2_, reducing safety concerns; and allow operation in an ambient atmosphere, potentially dramatically reducing operating costs as well. Furthermore, there is no obvious penalty for operating small-scale systems, meaning that EET-EMP could enable highly distributed energy storage. However, as of today there is no easily genetically-engineered microbe capable of both electron uptake by EET and CO_2_ fixation, meaning that this would need to be created. It is unclear if the reductions in cost and system complexity are worth the trade-off in the amount of complex microbe engineering that would be needed for such a feat. As of today, we are unaware of the full complement of genes needed for the reverse electron transport chain. Furthermore, it is unclear how easy it would be for self-assembly of the large area ECMs that this approach would rely upon. For ECMs with conductivities similar to those produced by *G. sulfurreducens* and *S. oneidensis* several square meters of ECM would be required for every square meter of solar panel. In the lab, ECMs with areas exceeding only a few square centimeters are rarely seen [69]. If the very high reported conductivities of cable bacteria ECMs can be reproduced, these could reduce the ECM area to only a few square centimeters. Recent developments in the construction of engineered biofilms [70] suggests that it might be possible to build a biologically synthesized conductive matrix that is tailored for electrosynthesis with low resistivity, high thickness, high area, and high accessibility for CO_2_ and product egress.

Recent developments in coupling photo-chemistry with EET [71] opens up the possibility of constructing quantum-dot (QD)-microbe hybrids that directly inject electrons in to the EET complex and then into metabolism. This would allow for the development of a system free of photovoltaics and electrodes that could be deployed at potentially extremely low cost. The possibility of adjusting the redox potential of the Mtr EET complex without significantly reducing efficiency (**Fig. S5**), along with the tunability of the electronic structure of quantum dots could allow significant room for engineering. Here, the potential for significant cost reduction could make for a significant payoff for the complex genetic engineering required to combine EET and carbon fixation.

The upper limits of efficiency of the EMP schemes presented here exceed those of all known forms of photosynthesis. Are these gains in efficiency worth pursuing? Can EMP achieve a significantly higher fraction of its theoretical efficiency in the real world than photosynthesis at an affordable cost? We cannot guarantee this, but the framework developed here gives us and other investigators the ability to rapidly understand the potential bang for buck of EMP schemes (of which there are many more than presented here). We hope that with the roadmap this framework gives, we and others in parallel can rapidly advance the field in multiple directions.

## Materials and Methods

The theory presented in this work was implemented in the RewiredCarbon suite of software developed with Python with the SciPy [72] and NumPy [73] libraries. Initial visualization was implemented with Matplotlib [74]. All computer code is available at github.com/barstowlab/rewiredcarbon

## Acknowledgment

This work was supported by a Career Award at the Scientific Interface from the Burroughs-Wellcome Fund (to B.B.), Princeton University startup funds (B.B.), Cornell University startup funds (B.B.), a Cornell Energy Systems Institute (CESI) Postdoctoral Fellowship (A.M.S) and by U.S. Department of Energy Biological and Environmental Research grant DE-SC0020179 (B.B.)

