## Supplementary material for "Constraints on the Efficiency of Electromicrobial Production": Compiled Supplementary Information (from another version of this preprint: version 1, 2020-06-24)

### Supplementary Information Contents For:
### Constraints on the Efficiency of Electromicrobial Production

*Farshid Salimijazi[a], Jaehwan Kim[b,c], Alexa M. Schmitz[a], Richard Grenville[d], Andrew Bocarsly[b], and Buz Barstow[a,1]*

*[a]Department of Biological and Environmental Engineering, Cornell University, Ithaca, NY 14853, USA;*
*[b]Department of Chemistry, Princeton University, Princeton, NJ 08540, USA;*
*[c]Present Address: Department of Chemistry and Chemical Biology, Cornell University, Ithaca, NY 14853, USA;*
*[d]Philadelphia Mixing Solutions Ltd., Palmyra, PA 17078, USA*

### Supplementary Information Contents

#### Supplementary Information Text


1. General Definition of Efficiency of *in vivo* $CO_2$-fixation Electromicrobial Production
2. Electron Requirements for Fuel Synthesis for *in vivo* $CO_2$-fixation $H_2$-mediated Electromicrobial Production
3. Total Cell and System Volume Requirements for Electromicrobial Production
4. $H_2$-transport by Diffusion
5. Cell Densities in Electromicrobial Production Systems
6. $H_2$-transport by Agitation
7. Electron Requirements for Fuel Synthesis for *in vivo* $CO_2$-fixation EET-mediated Electromicrobial Production
8. Calculation of Biofilm Overpotential and Resulting Electrical to Fuel Efficiency
9. Representative Biofilm Resistivities and Thicknesses
10. Efficiency of Electromicrobial Production with *ex vivo* Electrochemical $CO_2$-fixation
11. Metabolic Pathway Parameters


#### Supplementary Information Figures


1. Increasing transmembrane voltage reduces overall efficiency for $H_2$- and EET-mediated electromicrobial production schemes.
2. Metabolic pathways for isopropanol and butanol production.
3. Effect of varying NAD(P)H, Ferredoxin and ATP and requirements for electricity to butanol conversion efficiency.
4. Volumetric transfer rate ($k_L a$) of $H_2$ rises with increasing cell density and decreasing volume.
5. Effect of changing the redox potential of components of the Mtr complex on electrical to fuel efficiency.
6. Maximum height of cell culture that can be supplied with $H_2$ by diffusion and corresponding area of culture needed to convert 330 W of electrical power to butanol.
7. Maximum biofilm thickness and minimum area needed to achieve 50, 75 and 95% of peak efficiency.


#### Supplementary Information Tables


1. Symbols used in this article.
2. Metabolic pathway parameters
3. Representative extracellular matrix thickness ranges
4. Representative extracellular matrix resistivity ranges
5. Representative cell density ranges

### Supplementary Information Text for Constraints on the Efficiency of Electromicrobial Production

Farshid Salimijazi[1], Jaehwan Kim[2], Alexa Schmitz[1], Richard Grenville[3], Andrew Bocarsly[2], and Buz Barstow*[1]

[1]*Department of Biological and Environmental Engineering, Cornell University, Ithaca, NY 14853, USA*
[2]*Department of Chemistry, Princeton University, Princeton, NJ 08540, USA*
[3]*Philadelphia Mixing Solutions Ltd., Palmyra, PA 17078, USA*

#### 1 General Definition of Efficiency of *in vivo* $CO_2$-fixation Electromicrobial Production

In this work, we define the energy conversion efficiency of a electromicrobial production system as the rate of energy storage molecule production by the system, $\dot{N}_{\text{fuel}}$, multiplied by the energy content per molecule, $E_{\text{fuel}}$, relative to the total power input. **Figs. 1A** and **1B** in the main text show simplified schematics of electromicrobial production systems with *in vivo* enzymatic and *ex vivo* $CO_2$-fixation. In **Fig. 1A** a microbe absorbs electricity by $H_2$-oxidation or EET to generate reducing equivalents needed to enzymatically fix $CO_2$ and synthesize an energy storage molecule like polyhydroxybutyrate (PHB) or a hydrocarbon fuel at a rate $\dot{N}_{\text{fuel}}$. In **Fig. 1B** $CO_2$ is first electrochemically fixed to a lightly-reduced short chain hydrocarbon like formate or formic acid [1–3]. A microbe in the second cell absorbs electricity and further reduces and concatenates the initial fixation product a longer-chain hydrocarbon. This section deals with the efficiency of systems that use *in vivo* enzymatic $CO_2$-fixation. The efficiency of systems using *ex vivo* electrochemical $CO_2$-fixation is covered in **SI Text 10**.

The solar power to storage molecule, solar to electricity, and electricity to storage molecule conversion efficiencies,

$$\eta_{\text{SF}} = \dot{N}_{\text{fuel}}\, E_{\text{fuel}}/P_{\gamma}, \tag{1}$$

$$P_{\text{e, total}} = \eta_{\text{SE}}\, P_{\gamma}, \tag{2}$$

$$\eta_{\text{EF}} = \dot{N}_{\text{fuel}}\, E_{\text{fuel}}/P_{\text{e, total}}. \tag{3}$$

The input power to the electrochemical cell is the total electrical power, $P_{\text{e, total}}$, minus any power needed to agitate the system,

$$P_{\text{e, avail}} = P_{\text{e, total}} - P_{\text{e, stir}}. \tag{4}$$

*Corresponding author

Estimating the efficiency of *in vivo* $CO_2$-fixation electromicrobial production schemes (**Fig. 1A**) comes down to estimating $\dot{N}_{\mathrm{fuel}}$ as a function of the current flowing into the electrochemical system, $I_{\mathrm{cell}}$, and the voltage across the electrochemical cell, $\Delta U_{\mathrm{cell}}$,

$$\Delta U_{\mathrm{cell}} = U_{\mathrm{redox}} + U_{\mathrm{bias}}, \tag{5}$$
$$\Delta U_{\mathrm{cell}}\, I_{\mathrm{cell}} = P_{\mathrm{e,\ avail}}, \tag{6}$$

and the charge required to produce a single fuel molecule, $e\,\nu_{\mathrm{ef}}$ (the elementary charge multiplied by the number of electrons needed to synthesize a single fuel molecule),

$$\dot{N}_{\mathrm{fuel}} \leq I_{\mathrm{cell}}/e\,\nu_{\mathrm{ef}}, \tag{7}$$
$$\dot{N}_{\mathrm{fuel}} \leq P_{\mathrm{e,\ avail}}/\left(e\,\nu_{\mathrm{ef}}\,\Delta U_{\mathrm{cell}}\right). \tag{8}$$

Therefore, the overall electrical to fuel efficiency,

$$\eta_{\mathrm{EF}} \leq E_{\mathrm{fuel}}/\left(e\,\nu_{\mathrm{ef}}\,\Delta U_{\mathrm{cell}}\right). \tag{9}$$

### 2 Electron Requirements for Fuel Synthesis for *in vivo* $CO_2$-fixation $H_2$-mediated Electromicrobial Production

Estimating the efficiency of *in vivo* $CO_2$-fixation electromicrobial production systems (**Fig. 1A**) in the main text) comes down to estimating the total fuel production rate, $\dot{N}_{\mathrm{fuel}}$, from the electrical power available for electrochemistry, $P_{\mathrm{e,\ avail}}$, the voltage across the electrochemical cell, $\Delta U_{\mathrm{cell}}$, and the number of electrons needed to generate the NAD(P)H, Ferredoxin (Fd), and ATP for synthesis of a single fuel molecule from $CO_2$, $\nu_{\mathrm{ef}}$.

The number of electrons needed to synthesize a fuel molecule, $\nu_{\mathrm{ef}}$, can be estimated from molecular models of electron uptake. A schematic of the *Ralstonia eutropha* $H_2$-oxidation machinery (used by references [4–6]) is shown in **Fig. 2A** in the main text.

The low redox potential of $H_2$ ($U_{H_2}$) enables direct reduction of NADH by the cytosolic nickel-iron Soluble Hydrogenase (SH) (*R. eutropha* uses NADH rather than NADPH for $CO_2$-fixation) [7, 8]. While the *R. eutropha* genome does not code for any Ferredoxin-reducing di-iron hydrogenases, these could be readily added to it [9–11]. As $H_2$, NAD(P)H and Ferredoxin (Fd) all carry two electrons each, the microbe simply has to oxidize a number of $H_2$ molecules equal to the sum of NAD(P)H and Fd molecules that it needs to synthesize a fuel molecule from $CO_2$. In this work we consider only one $CO_2$-fixation cycle, the dicarboxylate/4-hydroxybutyrate (4HB) cycle [12], that requires Fd.

ATP is generated by injection of electrons from $H_2$-oxidation by the Membrane-Bound Hydrogenase (MBH) into the inner membrane electron transport chain [7,8]; proton pumping against the transmembrane voltage, $\Delta U_{\mathrm{membrane}}$; reduction of a terminal electron acceptor at a redox potential $U_{\mathrm{Acceptor}}$; and finally proton release through the ATP synthase.

The total minimum number of electrons needed to make a single fuel molecule from $CO_2$ using $H_2$ as an electron donor is just the sum of the electrons needed to reduce the NAD(P)H and Fd for the fuel molecule, $\nu_{\mathrm{ef,\,NADH}}$ and $\nu_{\mathrm{ef,\,Fd}}$, and the number of electrons that need to be sent downhill to make the ATP for the fuel molecule, $\nu_{\mathrm{ef,\,ATP,\,H_2}}$,

$$\nu_{\mathrm{ef,\,H_2}} = \nu_{\mathrm{ef,\,NADH}} + \nu_{\mathrm{ef,\,Fd}} + \nu_{\mathrm{ef,\,ATP,\,H_2}}. \tag{10}$$

As $H_2$, NAD(P)H and Ferredoxin (Fd) all carry two electrons each, the number of electrons needed to reduce them is just double the number of these molecules needed for fuel synthesis ($\nu_{\mathrm{f,\,NADH}}$ and $\nu_{\mathrm{f,\,Fd}}$),

$$\nu_{\mathrm{ef,\,H_2}} = 2\,\nu_{\mathrm{f,\,NADH}} + 2\,\nu_{\mathrm{f,\,Fd}} + \nu_{\mathrm{ef,\,ATP,\,H_2}}. \tag{11}$$

Estimates for the NAD(P)H, Fd and ATP requirements for isopropanol and 1-butanol synthesis (the metabolic pathways for synthesis of these fuels are shown in **Fig.S2** from $CO_2$ fixed by the known natural $CO_2$-fixation cycles and the synthetic CETCH cycle [13] are shown in **Supplementary Table 2**.

In the absence of specific metabolic pathway data, the absolute minimum value of $\nu_{\mathrm{ef,\,NADH}}$ can be calculated from the number of C-H, C-C and C-O bonds that need to be reduced during synthesis of the fuel from $CO_2$. For propanol ($C_3H_7OH$) this is 18, while for butanol ($C_4H_9OH$), it is 24. The number of NAD(P)H molecules, $\nu_{\mathrm{f,\,NADH}}$, is just half the number of electrons.

Likewise, we can estimate the number of ATP molecules needed to synthesize a single fuel molecule, $\nu_{\mathrm{f,\,ATP}}$, by looking to the ratio of ATP to NAD(P)H molecules consumed in naturally-occurring $CO_2$-fixation pathways and cycles. The ATP:NAD(P)H ratio varies from 0.25 in the production of acetyl-CoA in the Wood-Ljungdahl pathway to 1.5 in the production of glyceraldehyde-3-phosphate in the Calvin cycle. Similarly, in cyanobacterial photosynthesis for each $H_2O$ molecule that is split and NADPH molecule that is reduced, 6 protons are pumped across the thylakoid membrane (processes such as cyclic photophosphorylation can increase this number) [14], which is sufficient to catalyze the production of 4 ATP molecules [15]. Therefore, we assume at least 1.5 ATP molecules are required for each NAD(P)H molecule used in $CO_2$-fixation and subsequent synthesis of a biofuel. Thus, in the case of the synthesis of butanol from $CO_2$, 18 ATPs ($1.5 \times 12$) are required to produce one molecule of butanol.

In $H_2$-mediated electromicrobial production the cell current is used to reduce protons to $H_2$, generating an effective current, measured in Amps, that quantifies the number of electrons carried per second by $H_2$,

$$I_{\mathrm{H_2}} = I_{\mathrm{cell}}\,\xi_{\mathrm{eH_2}}, \tag{12}$$

where, for an upper estimate on efficiency, the Faraday efficiency of $H_2$-reduction $\xi_{\mathrm{eH_2}} = 1.0$.

The $H_2$-current is split into three (in most cases two) streams by the $H_2$-oxidizing microbe (**Fig. 2** in the main text). Two of the streams are used to directly produce the NAD(P)H and Fd used for $CO_2$-fixation. The third stream is injected into the electron transport chain where it flows downhill in energy to a terminal acceptor at a redox potential of $U_{\mathrm{Acceptor}}$ (usually $O_2$), building a proton gradient that is used to produce ATP,

$$I_{\mathrm{H_2}} = I_{\mathrm{NADH}} + I_{\mathrm{Fd}} + I_{\mathrm{downhill,\,ATP}}. \tag{13}$$

The maximum number of protons that can pumped from the cytosol to the periplasm against the transmembrane voltage $\Delta U_{\mathrm{membrane}}$, by sending one electron downhill to the terminal electron acceptor (floor rounds down to the nearest integer),

$$p_{\text{out, H}_2} = \text{floor}\left(\frac{U_{\text{H}_2} - U_{\text{Acceptor}}}{\Delta U_{\text{membrane}}}\right). \tag{14}$$

The minimum number of protons needed to be pumped from the periplasm back to the cytosol to produce a single ATP molecule (ceil rounds up to the nearest integer),

$$p_{\text{in, ATP}} = \text{ceil}\left(\frac{\Delta G_{\text{ATP/ADP}}}{e\,\Delta U_{\text{membrane}}}\right). \tag{15}$$

Thus, the number of electrons that need to be sent downhill to the terminal electron acceptor to make the ATP for a single fuel molecule,

$$\nu_{\text{ef, ATP, H}_2} = \frac{\nu_{\text{f, ATP}}\, p_{\text{in, ATP}}}{p_{\text{out, H}_2}}. \tag{16}$$

Thus, the total number of electrons that need to be sourced from $H_2$-oxidation to synthesize a single fuel molecule,

$$\begin{aligned} \nu_{\text{ef, H}_2} &= \nu_{\text{ef, NADH}} + \nu_{\text{ef, Fd}} + \nu_{\text{f, ATP}} \frac{\text{ceil}\left(\Delta G_{\text{ATP/ADP}}/e\,\Delta U_{\text{membrane}}\right)}{\text{floor}\left(\left(U_{\text{H}_2} - U_{\text{Acceptor}}\right)/\Delta U_{\text{membrane}}\right)}, & (17) \\ &= 2\,\nu_{\text{f, NADH}} + 2\,\nu_{\text{f, Fd}} + \nu_{\text{f, ATP}} \frac{\text{ceil}\left(\Delta G_{\text{ATP/ADP}}/e\,\Delta U_{\text{membrane}}\right)}{\text{floor}\left(\left(U_{\text{H}_2} - U_{\text{Acceptor}}\right)/\Delta U_{\text{membrane}}\right)}. & (18) \end{aligned}$$

Consequently, the electrical to fuel conversion efficiency for a $H_2$-mediated electromicrobial production scheme,

$$\eta_{\text{EH}_2\text{F}} = \frac{E_{\text{fuel}}}{e\,\Delta U_{\text{cell}}\,\nu_{\text{ef, H}_2}}, \tag{19}$$

and the solar to electricity to fuel conversion efficiency,

$$\eta_{\text{SH}_2\text{F}} = \frac{\eta_{\text{SE}}\,E_{\text{fuel}}}{e\,\Delta U_{\text{cell}}\,\nu_{\text{ef, H}_2}}, \tag{20}$$

where $\eta_{\text{SE}}$ is the solar to electrical energy conversion efficiency [16].

### 3 Total Cell and System Volume Requirements for Electromicrobial Production

How many electroactive microbes are needed to store a given amount of renewable electricity? We first considered the answer to this question in an earlier article [8], but restate and refine these arguments here for convenience, and with a set of symbols consistent with those in the rest of this article.

In most scenarios examined in this article, we consider the current, $I_\gamma$, and voltage, $U_\gamma$, produced by an idealized $1\,\text{m}^2$ single-junction Si solar photovoltaic illuminated by the $1{,}000\,\text{W}\,\text{m}^{-2}$ AM1.5

reference spectrum [17, 18] operating at its peak power point of 0.75 V and 430 A [19]. Unless otherwise noted, all calculations assume ambient temperature (300 K).

To operate a bio-electrochemical cell, the photo-current and photo-voltage are transformed by a potentiostat that is assumed to be perfectly efficient (we are making an upper limit estimate of efficiency). By conservation of energy, the electrical power from the solar PV will be equal to the electrical power in the bio-electrochemical cell,

$$I_\gamma \, U_\gamma = I_{\text{cell}} \, \Delta U_{\text{cell}}. \tag{21}$$

We refer to the effective current available to biological process as $I_{\text{cell, bio}}$.

In $H_2$-mediated electromicrobial production systems, the cell current is used to reduce protons to $H_2$, generating an effective current, measured in Amps, that quantifies the number of electrons carried per second by $H_2$ that is available for $H_2$-oxidizing microbes,

$$I_{\text{cell, bio}} = I_{\text{H}_2}, \tag{22}$$

$$I_{\text{H}_2} = I_{\text{cell}} \, \xi_{\text{eH}_2}, \tag{23}$$

where, for an upper estimate on efficiency, the Faraday efficiency of $H_2$-reduction $\xi_{\text{eH}_2} = 1.0$.

In extracellular electron transfer (EET)-mediated systems, the cell current is used to reduce the Mtr complex that connects the cell surface to the electron transport chain on the inner membrane, which is then used to reduce the quinone pool. This generates effective currents $I_{\text{Mtr}}$ and $I_{\text{Q}}$.

$$I_{\text{Mtr}} = \xi_{\text{eMtr}} \, I_{\text{cell}}, \tag{24}$$

$$I_{\text{Q}} = \xi_{\text{eMtrQ}} \, I_{\text{Mtr}}, \tag{25}$$

$$I_{\text{Q}} = \xi_{\text{eMtr}} \, \xi_{\text{eMtrQ}} \, I_{\text{cell}} \tag{26}$$

We assume, (as we are calculating an upper bound on efficiency) that all of the cell current going into the cell ($I_{\text{cell}}$), is transferred to Mtr complex ($I_{\text{Mtr}}$), and from there, to the quinone pool ($I_{\text{Q}}$) with no charge losses ($\xi_{\text{eMtr}} = \xi_{\text{eMtrQ}} = 1$). Thus,

$$I_{\text{Q}} = I_{\text{cell, bio}} \tag{27}$$

As the $CO_2$ fixation rate is limiting [8], the number of electrons consumed by each cell per unit time is a function of the number of electrons needed to synthesize each fuel molecule, $\nu_{\text{ef}}$ (calculated in **SI Text 2** for $H_2$-mediated systems and in **SI Text 7** for EET-mediated systems), the number of carbon atoms needed to be fixed per fuel molecule (not the number incorporated which is often less) $\nu_{\text{Cf, fix}}$ and the rate and number of carbon-fixing enzymes, $r_{\text{fix}}$ and $\nu_{\text{fix}}$. The electron consumption rate per cell,

$$\dot{\nu}_e = \frac{\nu_{\text{ef}} \, r_{\text{fix}} \, \nu_{\text{fix}}}{\nu_{\text{Cf, fix}}}. \tag{28}$$

Thus, the total number of cells needed to deal with the solar power is,

$$N_{\text{cells}} = \frac{I_{\text{cell, bio}}}{e \, \dot{\nu}_e}, \tag{29}$$

and the volume of cells,

$$V_{\text{cells}} = N_{\text{cells}} / \text{n}_{\text{cells}}. \tag{30}$$

#### 4 $H_2$-transport by Diffusion

A commonly expressed concern regarding $H_2$-mediated electromicrobial production systems is the limitation on $H_2$ transport due to its low solubility in water. We first considered the geometric constraints imposed on a electromicrobial production system by the low solubility of $H_2$ in an earlier article [8], but restate and refine the arguments here with a set of symbols consistent with those in the rest of this article.

Consider a volume of cell culture, $V_{\text{cells}}$, containing cells at a density, $\text{n}_{\text{cells}}$. $H_2$ gas at a pressure $P_{H_2}$ is supplied to these cells through the top surface of this volume, which has an area $A_{\text{cell}}$. Each cell consumes electrons at a rate $\dot{\nu}_e$ in order to fix $CO_2$ and synthesize an energy storage molecule (this rate is calculated in **SI Text 2**). The total $H_2$ demand of these cells (given that $H_2$ carries 2 electrons per molecule) is,

$$\dot{\nu}_{H_2} = \frac{1}{2}\,\dot{\nu}_e\,\text{n}_{\text{cells}}\,V_{\text{cells}}. \tag{31}$$

Fick's first law can be used to calculate the size of the $H_2$ concentration, $C_{H_2}$, gradient necessary to produce this flux [20]. The rate of diffusional particle flux into the volume $V_{\text{cells}}$ through a planar face of area $A_{\text{cell}}$ is equal to the concentration gradient normal to that plane multiplied by the diffusion coefficient of the particle species,

$$\frac{1}{A_{\text{film}}}\frac{dN_{H_2}}{dt} = -D_{H_2}\frac{dC_{H_2}}{dx}, \tag{32}$$

$$\frac{1}{A_{\text{film}}}\frac{\dot{\nu}_{H_2}}{N_{\text{A}}} = -D_{H_2}\frac{dC_{H_2}}{dx}. \tag{33}$$

The cell culture volume is the product of the height of the culture, $h_{\text{film}}$, and the area of the cross section,

$$V_{\text{cells}} = A_{\text{film}}\,h_{\text{film}}. \tag{34}$$

The $H_2$ concentration at the top surface of the cell culture is assumed to be equal to the maximum solubility of $H_2$ under the conditions of the system. This can be calculated with Henry's Law, as a function of the partial pressure of $H_2$ in the headspace, $P_{H_2}$ and the solubility constant for $H_2$ in water, $k_{H_2}$,

$$C_{H_2,\,\text{max}} = P_{H_2}/k_{H_2}. \tag{35}$$

Over the height of the culture volume, $h_{\text{film}}$, the $H_2$ concentration drops to 0 due to consumption by the cells,

$$\frac{1}{A_{\text{film}}}\frac{\dot{\nu}_{H_2}}{N_{\text{A}}} = D_{H_2}\frac{C_{H_2,\,\text{max}}}{h_{\text{film}}}. \tag{36}$$

Thus,

$$
\begin{aligned}
h_{\mathrm{film}} &= \frac{D_{\mathrm{H_2}}\,N_{\mathrm{A}}\,A_{\mathrm{film}}\,C_{\mathrm{H_2,\ max}}}{\dot{\nu}_{\mathrm{H_2}}}, && (37)\\
h_{\mathrm{film}} &= \frac{D_{\mathrm{H_2}}\,N_{\mathrm{A}}\,A_{\mathrm{film}}\,P_{\mathrm{H_2}}}{k_{\mathrm{H_2}}\dot{\nu}_{\mathrm{H_2}}}, && (38)\\
h_{\mathrm{film}} &= \frac{2D_{\mathrm{H_2}}\,N_{\mathrm{A}}\,A_{\mathrm{film}}\,P_{\mathrm{H_2}}}{k_{\mathrm{H_2}}\,\mathrm{n_{cells}}\,\dot{\nu}_e\,V_{\mathrm{cells}}}, && (39)\\
h_{\mathrm{film}} &= \frac{2D_{\mathrm{H_2}}\,N_{\mathrm{A}}\,A_{\mathrm{film}}\,P_{\mathrm{H_2}}}{k_{\mathrm{H_2}}\,\mathrm{n_{cells}}\,\dot{\nu}_e\,A_{\mathrm{film}}\,h_{\mathrm{film}}}. && (40)
\end{aligned}
$$

Thus, the height of the film of cell culture,

$$
h_{\mathrm{film}} = \sqrt{(2P_{\mathrm{H_2}}\,D_{\mathrm{H_2}}\,N_{\mathrm{A}})/(k_{\mathrm{H_2}}\,\mathrm{n_{cells}}\,\dot{\nu}_e)}, \qquad (41)
$$

and its cross-sectional area,

$$
\begin{aligned}
A_{\mathrm{film}} &= V_{\mathrm{cells}}/h_{\mathrm{film}}, && (42)\\
A_{\mathrm{film}} &= \frac{I_{\mathrm{H_2}}}{e\,\dot{\nu}_e\,\mathrm{n_{cells}}\,h_{\mathrm{film}}}, && (43)\\
A_{\mathrm{film}} &= \frac{\xi_{\mathrm{eH_2}} I_{\mathrm{cell}}}{e\,\dot{\nu}_e\,\mathrm{n_{cells}}\,h_{\mathrm{film}}}, && (44)\\
A_{\mathrm{film}} &= \frac{\xi_{\mathrm{eH_2}} P_{\mathrm{e,\ avail}}}{e\,\dot{\nu}_e\,\mathrm{n_{cells}}\,h_{\mathrm{film}}\,\Delta U_{\mathrm{cell}}}. && (45)
\end{aligned}
$$

Assuming that no power is diverted to agitation, and that the electromicrobial production system is supplied by a solar photovoltaic with a solar to electrical efficiency $\eta_{\mathrm{SE}}$, illuminated by total solar power $P_\gamma$,

$$
\begin{aligned}
P_{\mathrm{e,\ avail}} &= P_{\mathrm{e,\ total}}, && (46)\\
P_{\mathrm{e,\ avail}} &= \eta_{\mathrm{SE}}\,P_\gamma. && (47)
\end{aligned}
$$

Assuming that the solar power incident upon the solar photovoltaic is the photovoltaic area, $A_{\mathrm{PV}}$, multiplied by the solar constant of $\approx 1{,}000\,\mathrm{W\,m^{-2}}$, $G_{\mathrm{SC}}$,

$$
P_\gamma = A_{\mathrm{PV}}\,G_{\mathrm{SC}}. \qquad (48)
$$

Thus, the area of the film of cell culture,

$$
\begin{aligned}
A_{\mathrm{film}} &= \frac{\xi_{\mathrm{eH_2}}\,\eta_{\mathrm{SE}}\,A_{\mathrm{PV}}\,G_{\mathrm{SC}}}{e\,\dot{\nu}_e\,\mathrm{n_{cells}}\,h_{\mathrm{film}}\,\Delta U_{\mathrm{cell}}}, && (49)\\
A_{\mathrm{film}} &= \frac{\xi_{\mathrm{eH_2}}\,\eta_{\mathrm{SE}}\,P_\gamma\,k_{\mathrm{H_2}}^{1/2}}{e\,\Delta U_{\mathrm{cell}}\,\left(2\dot{\nu}_e\,\mathrm{n_{cells}}\,P_{\mathrm{H_2}}\,D_{\mathrm{H_2}}\,N_{\mathrm{A}}\right)^{1/2}}, && (50)\\
A_{\mathrm{film}} &= \frac{\xi_{\mathrm{eH_2}}\,k_{\mathrm{H_2}}^{1/2}\,P_{\mathrm{e,\ avail}}}{e\,\Delta U_{\mathrm{cell}}\,\left(2\dot{\nu}_e\,\mathrm{n_{cells}}\,P_{\mathrm{H_2}}\,D_{\mathrm{H_2}}\,N_{\mathrm{A}}\right)^{1/2}}. && (51)
\end{aligned}
$$

### 5 Cell Densities in Electromicrobial Production Systems

What sorts of cell densities could we expect in a electromicrobial production system? We consider 4 cell density ranges: (**1**) laboratory grown cultures of *E. coli* in exponential phase; (**2**) cyanobacteria grown to maximum density; (**3**) cultures of *E. coli* at saturating density; (**4**) $H_2$-oxidizing microbes grown to maximum density; and (**5**) and saturating cultures of industrially-grown yeast.

**Density Range 1,** $\rho_1$. Laboratory grown cultures of *E. coli* with an optical density of 0.1 in LB-broth have number densities of between $2 \times 10^{13}$ (strain MG1655, BNID 103625) and $1 \times 10^{14}$ cells $m^{-3}$ (BNID 100985).

**Density Range 2,** $\rho_2$. What about cells grown under autotrophic conditions? Cyanobacteria typically have cell densities of $1\,g\,L^{-1}$ of cell dry weight. Assuming a cell mass of 148 to 280 fg, this corresponds to a density of $3.6 \times 10^{15}$ to $6.8 \times 10^{15}$ cells $m^{-3}$ [21]. Lippi *et al.* [21] report higher cyanobacterial densities of $\approx 30\,g\,L^{-1}$ of cell dry weight, but we consider these atypical.

**Density Range 3,** $\rho_3$. *E. coli* grown on LB-broth has a maximum culture density of 1.6 to $20.5\,g\,L^{-1}$ (BioNumbers ID (BNID) 104940). Assuming a dry weight of *E. coli* of 148 to 211 fg (BNID 102231) to 280 fg per cell (BNID 100008 and 103904), this corresponds to a cell number density of $5.1 \times 10^{15}\,m^{-3}$ to $1.39 \times 10^{17}$ cells $m^{-3}$.

**Density Range 4,** $\rho_4$. What about $H_2$-oxidizing cells grown under autotrophic conditions. The most relevant number is cell mass density reported by Tanaka *et al.* for *Ralstonia eutropha* (referred to at the time as *Alcaligenes eutrophus*) of $9.13 \times 10^4\,g\,m^{-3}$ [22]. Of this, the biopolymer polyhydroxybutyrate (PHB) accounts for $6.19 \times 10^4\,g\,m^{-3}$. While there is little data on the dry weight on an individual *R. eutropha* cell with or without PHB, assuming a similar weight as *E. coli* for the PHB-free *R. eutropha* cell, this corresponds to a cell number density of $1.0 \times 10^{17}$ to $2.0 \times 10^{17}$ cells $m^{-3}$.

**Density Range 5,** $\rho_5$. Industrially-grown strains of yeast have densities of between 10 to $150\,g\,L^{-1}$ [23]. Assuming a density of $10^{13}$ yeast cells $g^{-1}$ of dry weight, this corresponds to a number density of $1 \times 10^{17}$ to $1.5 \times 10^{18}$ cells $m^{-3}$. We consider $1.5 \times 10^{18}$ cells $m^{-3}$ to be an upper limit on cell density in industrial processes. Thus, it is reasonable to an upper limit on cell density of $\approx 1.5 \times 10^{18}$ cells $m^{-3}$ in a electromicrobial production system.

This discussion is summarized in **Table S5**.

### 6 $H_2$-transport by Agitation

What power is required to transport continuously produced $H_2$ into a continuously $H_2$-oxidizing cell culture by agitation? To determine this we iteratively solved a set of equations linking $H_2$ production, consumption, and the power required for gas mixing using a formalism compiled by Van't Riet [24] through the `Efficiency_Hydrogen_BioCO2_ to_Fuel_ScaleUp_Stirring_Fixed_CellDensity_ Grenville` algorithm in the REWIREDCARBON package until a self consistent set of solutions were found.

The total electrical power supplied to the electromicrobial production system is divided between the electrochemical cell that produces $H_2$, and an agitator that mixes the cell culture to distribute $H_2$ allowing it to be oxidized,

$$P_{\text{e, total}} = P_{\text{e, avail}} + P_{\text{e, stir}}. \tag{52}$$

The power flowing to the $H_2$-producing electrochemical cell drives a current, $I_{\text{cell}}$, across a potential difference $\Delta U_{\text{cell}}$, that is used to reduce protons to $H_2$, generating an effective current, $I_{\text{H}_2}$, measured in Amps, that quantifies the number of electrons carried per second by $H_2$ (**Equation S23**),

$$I_{\text{H}_2} = I_{\text{cell}}\, \xi_{\text{eH}_2}, \tag{53}$$
$$I_{\text{H}_2} = \xi_{\text{eH}_2} P_{\text{e, avail}} / \Delta U_{\text{cell}}. \tag{54}$$

Initially, it is assumed that $P_{\text{e, avail}} = P_{\text{e, total}}$.

In order for the $H_2$ fed into the tank to be continuously oxidized, the total $H_2$ uptake rate, set by the total number of cells in the culture, $N_{\text{cells}}$, the cell density, $\text{n}_{\text{cells}}$, and the electron uptake rate per cell, $\dot{\nu}_e$, (**SI Text 3**) must match the input rate,

$$I_{\text{H}_2} = e\, N_{\text{cells}}\, \dot{\nu}_e, \tag{55}$$
$$N_{\text{cells}} = I_{\text{H}_2} / e\, \dot{\nu}_e. \tag{56}$$

Thus, the volume of this cell culture,

$$V_{\text{cells}} = N_{\text{cells}} / \text{n}_{\text{cells}}. \tag{57}$$

The molar transfer rate of $H_2$ into the culture can be calculated from the $H_2$ current,

$$J_{\text{H}_2} = \frac{I_{\text{H}_2}}{2e\, N_{\text{A}}}, \tag{58}$$

while the volumetric transfer rate can be calculated with the ideal gas law from the $H_2$ current, the ambient temperature and the partial pressure of $H_2$ in the headspace of the tank,

$$\dot{v}_{\text{H}_2} = \frac{I_{\text{H}_2}\, k_{\text{B}}\, T}{2e\, P_{\text{H}_2}}. \tag{59}$$

We assume that the $H_2$ concentration at a bubble-liquid interface is the maximum allowed by Henry's law,

$$C_{\text{H}_2\text{, interface}} = \frac{P_{\text{H}_2}}{k_{\text{H}_2}}, \tag{60}$$

and that the bulk concentration is a fraction, $f_{\text{bulk to peak}}$, of this,

$$C_{\text{H}_2\text{, bulk}} = f_{\text{bulk to peak}}\, C_{\text{H}_2\text{, interface}}. \tag{61}$$

While $f_{\text{bulk to peak}}$ is a free parameter in REWIREDCARBON, for all calculations in this work we assume that $H_2$ consumption is so rapid that the $H_2$ concentration in the bulk liquid drops to zero,

$$f_{\text{bulk to peak}} = 0, \tag{62}$$
$$C_{\text{H}_2\text{, bulk}} = 0. \tag{63}$$

Thus, the difference in $H_2$ concentration, $\Delta C_{H_2}$, between the liquid side of the gas-liquid interface at the edge of a bubble and the gas concentration in the bulk liquid, $C_{H_2,\text{ bulk}}$,

$$\begin{aligned} \Delta C_{H_2} &= C_{H_2,\text{ interface}} - C_{H_2,\text{ bulk}}, && (64) \\ \Delta C_{H_2} &= P_{H_2}/k_{H_2}. && (65) \end{aligned}$$

We considered a system in which the cell culture is contained in a cylindrical tank with liquid height $h_{\text{tank}}$, a cross-sectional diameter $d_{\text{tank}}$, into which $H_2$ is introduced below the liquid surface, and is distributed by an agitator (**Fig. 5A**). The ratio of liquid height to tank diameter is $r_{\text{Dh}}$,

$$d_{\text{tank}} = r_{\text{Dh}}\, h_{\text{tank}}. \quad (66)$$

Thus,

$$\begin{aligned} V_{\text{cells}} &= \frac{\pi d_{\text{tank}}^3}{4 r_{\text{Dh}}}, && (67) \\ d_{\text{tank}} &= \left(4 r_{\text{Dh}} V_{\text{cells}}/\pi\right)^{1/3}, && (68) \end{aligned}$$

and the cross sectional area of the tank,

$$A_{\text{tank}} = \frac{\pi^{1/3}\left(V_{\text{cells}} r_{\text{Dh}}\right)^{2/3}}{4^{1/3}}. \quad (69)$$

While $r_{\text{Dh}}$ is a free parameter in REWIREDCARBON, for all calculations in this work we assumed that it was equal to 1.

The volume of gas passing through each unit area of the top surface of cell culture (the superficial gas velocity, $u_{\text{SG}}$),

$$u_{\text{SG}} = \dot{v}_{\text{gas}}/A_{\text{tank}}. \quad (70)$$

The volumetric mass transfer rate, $k_{\text{L}}a_{H_2}$, can be calculated from the molar transfer rate, $J_{H_2}$, and the difference in $H_2$ concentration between the gas-liquid interface at the edge of a bubble and the gas concentration in the bulk liquid, $\Delta C_{H_2}$,

$$k_{\text{L}}a_{H_2} = \frac{J_{H_2}}{V_{\text{cells}}\, \Delta C_{H_2}}. \quad (71)$$

The electrical power needed to stir $H_2$ into the cell culture can be estimated from an empirical formula that relates stirring power density, $\epsilon_{\text{gas}}$ [24],

$$P_{\text{e, stir}} = V_{\text{cells}}\, \epsilon_{\text{gas}}, \quad (72)$$

to the superficial gas velocity and volumetric mass transfer rate,

$$k_{\text{L}}a = A\, \epsilon_{\text{gas}}^B\, u_{\text{SG}}^C, \quad (73)$$

where the constants $A = 0.026$, $B = 0.4$, and $C = 0.5$ for aqueous solutions when the units of $k_{\text{L}}a_{H_2}$ are $\text{W m}^{-3}$, the units of $\dot{v}_{\text{gas}}$ are $\text{m}^3\,\text{s}^{-1}$, and the units of $u_{\text{SG}}$ are $\text{m s}^{-1}$ [24]. Thus,

$$\epsilon_{\text{gas}}^B = \frac{k_{\text{L}}a_{H_2}}{A\, u_{\text{SG}}^C}. \quad (74)$$

This can be rearranged

$$\epsilon_{\rm gas} = \left(\frac{k_{\rm L} a_{\rm H_2}}{A\, u_{\rm SG}^C}\right)^{1/B}, \quad (75)$$

$$P_{\rm e,\, stir} = V_{\rm cells} \left(\frac{k_{\rm L} a_{\rm H_2}}{A\, u_{\rm SG}^C}\right)^{1/B}. \quad (76)$$

To solve these equations, the initial guess for $P_{\rm e,\, stir}$ that was used to estimate $I_{\rm H_2}$ in **Equation S55** was increased in small increments until it reached the maximum available power, $P_{\rm e,\, total}$. The best guess for $P_{\rm e,\, stir}$ was the value that minimized the difference between the initial guess and the value calculated in **Equation S76**. This value was further refined with the SCIPY.OPTIMIZE package.

### 7 Electron Requirements for Fuel Synthesis for *in vivo* $CO_2$-fixation EET-mediated Electromicrobial Production

While $H_2$ has a sufficiently low redox potential ($U_{\rm H_2} = -0.42$ V relative to the Standard Hydrogen Electrode (SHE) at pH 7.0) to directly reduce NAD(P)$^+$ to NAD(P)H ($U_{\rm NADH} = -0.32$ V) and Ferredoxin (Fd, $U_{\rm Fd} \approx -0.4$ V [25]) (**Fig. 2** in the main text), the Mtr EET complex and its homologs built into electroactive microbes ($U_{\rm Mtr}$ has an average value of $\approx -0.2$ V vs. SHE [26]), and the quinone pool in the inner membrane ($U_{\rm Q} = -0.067$ to $-0.11$ V vs. SHE for menaquinone [27,28], or $+0.11$ V for ubiquinone [29]) with which it interacts do not (**Fig. 2**). This necessitates a mechanism, sometimes referred to as the "uphill pathway" or reverse electron transport, to raise the energy of electrons absorbed by the cell from high redox potential sources that is analogous to a transformer. However, unlike an inductive transformer used in electronic technology it is postulated that this mechanism operates electrochemically [30].

It is hypothesized that electroactive microbes capable of electron uptake from high redox potential sources like iron minerals transfer electrons from an external substrate through the EET complex that spans the outer and inner membranes and then onto the quinone pool in the inner membrane at a redox potential $U_{\rm Q}$ (**Fig. 2B**). Up until this point, the flow electrons is downhill in energy. However, the redox potential of menaquinone is $\approx 200$ mV above that of NAD(P)H, while ubiquinone is $\approx 400$ mV above it [30]. Somehow these electrons need to gain between 200 and 400 meV of energy in order to reduce NAD(P)H.

Furthermore, it is hypothesized that the reverse electron transport chain shunts some of the incoming electron flow downhill to a terminal electron acceptor such as $O_2$ at a redox potential $U_{\rm Acceptor}$, in the process pumping protons against the inner membrane potential difference ($\Delta U_{\rm membrane}$) and generating a proton gradient across the inner membrane. Some of the energy stored in the proton gradient is used to drive ATP production, while the remainder is used to provide an energy source for reduction of NAD(P)H.

We assume that the cell current, $I_{\rm cell}$, is transferred to the Mtr EET complex with a Faradaic efficiency $\xi_{\rm eMtr}$,

$$I_{\rm Mtr} = \xi_{\rm eMtr}\, I_{\rm cell}. \quad (77)$$

The current flowing through the Mtr complex is then transferred ot the quinone pool with an efficiency $\xi_{\text{eMtrQ}}$,

$$I_{\text{Q}} = \xi_{\text{eMtrQ}}\, I_{\text{Mtr}}. \tag{78}$$

Thus,

$$I_{\text{Q}} = \xi_{\text{eMtrQ}}\, \xi_{\text{eMtr}}\, I_{\text{cell}}. \tag{79}$$

We assume, (as we are calculating an upper bound on efficiency) that all of the cell current is transferred to Mtr complex and from there to the quinone pool with no charge losses ($\xi_{\text{eMtr}} = \xi_{\text{eMtrQ}} = 1$).

After reaching the quinone pool, the current flow is divided, with some used for direct NAD(P)H and Ferredoxin (Fd) reduction, and some sent downhill to generate the proton gradient needed to produce ATP as in the $H_2$-oxidation case. However, an additional current needs to be sent downhill to produce a proton gradient to raise the energy of electrons used for NAD(P)H and Fd reduction,

$$I_{\text{Q}} = I_{\text{NADH}} + I_{\text{Fd}} + I_{\text{downhill, ATP}} + I_{\text{downhill, NADH}} + I_{\text{downhill, Fd}}. \tag{80}$$

Following the same logic used to describe $H_2$-oxidation (**SI Text 2**), the total number of electrons needed to be acquired by EET in order to synthesize a single fuel molecule is the sum of the electrons needed to reduce the NAD(P)H and Fd for the fuel molecule, $\nu_{\text{ef, NADH}}$ and $\nu_{\text{ef, Fd}}$; the number of electrons that need to be sent downhill to regenerate the ATP for the fuel molecule, $\nu_{\text{ef, ATP, EET}}$; and finally the number of electrons needed to be sent downhill to provide energy for NAD(P)H and Fd reduction, $\nu_{\text{ef, NADH, down}}$ and $\nu_{\text{ef, Fd, down}}$. Thus,

$$\nu_{\text{ef, EET}} = \nu_{\text{ef, NADH}} + \nu_{\text{ef, Fd}} + \nu_{\text{ef, ATP, EET}} + \nu_{\text{ef, NADH, down}} + \nu_{\text{ef, Fd, down}}. \tag{81}$$

As in the $H_2$-oxidation case, the maximum number of protons that can pumped from the cytosol to the periplasm against the transmembrane voltage $\Delta U_{\text{membrane}}$, by sending one electron downhill to the terminal electron acceptor,

$$p_{\text{out, EET}} = \text{floor}\left(\frac{U_{\text{Q}} - U_{\text{Acceptor}}}{\Delta U_{\text{membrane}}}\right). \tag{82}$$

Note that $p_{\text{out, EET}}$ is smaller than $p_{\text{out, H}_2}$, as the starting redox potential of the electrons, $U_{\text{Q}}$, is higher than $U_{\text{H}_2}$.

As in the $H_2$-oxidation case, the number of protons that need to released through the ATP synthase in order to regenerate one molecule of ATP,

$$p_{\text{in, ATP}} = \text{ceil}\left(\frac{\Delta G_{\text{ATP/ADP}}}{e\,\Delta U_{\text{membrane}}}\right), \tag{83}$$

Continuing to follow this logic, the number of protons that need to be pumped from the periplasm to the cytosol in order to raise one electron to the energy of NAD(P)H,

$$p_{\text{in, NADH}} = \text{ceil}\left(\frac{U_{\text{NADH}} - U_{\text{Q}}}{\Delta U_{\text{membrane}}}\right), \tag{84}$$

and the number of protons that need to be pumped from the periplasm to the cytoplasm to raise on electron to the energy of Fd,

$$p_{\text{in, Fd}} = \text{ceil}\left(\frac{U_{\text{Fd}} - U_{\text{Q}}}{\Delta U_{\text{membrane}}}\right). \tag{85}$$

Thus, the number of electrons that need to be sent downhill to regenerate the ATP ($\nu_{\text{f, ATP}}$) needed to synthesize one fuel molecule,

$$\nu_{\text{ef, ATP, EET}} = \frac{\nu_{\text{f, ATP}}\, p_{\text{in, ATP}}}{p_{\text{out, EET}}}. \tag{86}$$

The total number of electrons that need to be sent downhill in order to raise $\nu_{\text{ef, NADH}}$ electrons in energy in order to reduce $\nu_{\text{f, NADH}}$ NAD(P)H molecules needed to synthesize a single fuel molecule,

$$\nu_{\text{ef, NADH, down}} = \frac{\nu_{\text{ef, NADH}}\, p_{\text{in, NADH}}}{p_{\text{out, EET}}}. \tag{87}$$

The total number of electrons that need to be sent downhill in order to raise $\nu_{\text{ef, Fd}}$ electrons in energy in order to reduce $\nu_{\text{f, Fd}}$ Fd molecules needed to synthesize a single fuel molecule,

$$\nu_{\text{ef, Fd, down}} = \frac{\nu_{\text{ef, Fd}}\, p_{\text{in, Fd}}}{p_{\text{out, EET}}}. \tag{88}$$

Thus, the total number of electrons that need to be acquired by EET to synthesize a single fuel molecule,

$$\begin{aligned} \nu_{\text{ef, EET}} = {} & \nu_{\text{ef, NADH}} + \nu_{\text{ef, Fd}} \\ & + \nu_{\text{f, ATP}} \frac{\text{ceil}\left(\Delta G_{\text{ATP/ADP}}/e\,\Delta U_{\text{membrane}}\right)}{\text{floor}\left(\left(U_{\text{Q}} - U_{\text{Acceptor}}\right)/\Delta U_{\text{membrane}}\right)} \\ & + \nu_{\text{ef, NADH}} \frac{\text{ceil}\left(U_{\text{NADH}} - U_{\text{Q}}/\Delta U_{\text{membrane}}\right)}{\text{floor}\left(\left(U_{\text{Q}} - U_{\text{Acceptor}}\right)/\Delta U_{\text{membrane}}\right)} \\ & + \nu_{\text{ef, Fd}} \frac{\text{ceil}\left(U_{\text{Fd}} - U_{\text{Q}}/\Delta U_{\text{membrane}}\right)}{\text{floor}\left(\left(U_{\text{Q}} - U_{\text{Acceptor}}\right)/\Delta U_{\text{membrane}}\right)}. \end{aligned} \tag{89}$$

As two electrons are required to reduce both NAD(P)H and Fd,

$$\begin{aligned} \nu_{\text{ef, EET}} = {} & 2\,\nu_{\text{f, NADH}} + 2\,\nu_{\text{f, Fd}} \\ & + \nu_{\text{f, ATP}} \frac{\text{ceil}\left(\Delta G_{\text{ATP/ADP}}/e\,\Delta U_{\text{membrane}}\right)}{\text{floor}\left(\left(U_{\text{Q}} - U_{\text{Acceptor}}\right)/\Delta U_{\text{membrane}}\right)} \\ & + 2\,\nu_{\text{f, NADH}} \frac{\text{ceil}\left(U_{\text{NADH}} - U_{\text{Q}}/\Delta U_{\text{membrane}}\right)}{\text{floor}\left(\left(U_{\text{Q}} - U_{\text{Acceptor}}\right)/\Delta U_{\text{membrane}}\right)} \\ & + 2\,\nu_{\text{f, Fd}} \frac{\text{ceil}\left(U_{\text{Fd}} - U_{\text{Q}}/\Delta U_{\text{membrane}}\right)}{\text{floor}\left(\left(U_{\text{Q}} - U_{\text{Acceptor}}\right)/\Delta U_{\text{membrane}}\right)}. \end{aligned} \tag{90}$$

Thus, the electrical energy to fuel efficiency of EET-mediated electromicrobial production is,

$$\eta_{\text{EEF}} = \frac{E_{\text{fuel}}}{e\,\Delta U_{\text{cell}}\, \nu_{\text{ef, EET}}} \tag{91}$$

### 8 Calculation of Biofilm Overpotential and Resulting Electrical to Fuel Efficiency

We calculate the efficiency of the system by calculating the overpotential of a biofilm. We solved for the cell current $I_{\text{cell}}$, the bias voltage needed to drive current across the biofilm $\Delta U_{\text{biofilm}}$, the area of the biofilm $A_{\text{biofilm}}$, the total number of cells in the biofilm $N_{\text{cells}}$, and the volume of the biofilm $V_{\text{biofilm}}$.

We also included the parameters, the thermodynamic minimum voltage across the cell $U_{\text{redox}}$ set by the half-potentials of Mtr and water splitting at the anode, any additional bias that needs to be applied across the cell $U_{\text{bias}}$, the electron uptake rate per cell $\dot{\nu}_e$, the thickness of the biofilm $h_{\text{biofilm}}$, the resistivity of the biofilm $\rho_{\text{biofilm}}$, the Faradaic efficiency of transfer of electrons from the electrode to Mtr $\xi_{\text{eMtr}}$, the Faradaic efficiency of transfer of electrons from Mtr to the quinone pool $\xi_{\text{eMtrQ}}$, the fraction of the area of each layer of cells occupied by cells, or fill factor $f_{\text{layer}}$, the height of a layer of cells in the biofilm $h_{\text{layer}}$, and the cross sectional area of a cell $A_{\text{cell}}$.

We developed five coupled equations,

$$I_{\text{cell}} = \frac{P_{\text{e, avail}}}{U_{\text{redox}} + U_{\text{bias}} + \Delta U_{\text{biofilm}}} \tag{92}$$

$$\Delta U_{\text{biofilm}} = \frac{\rho_{\text{biofilm}}\, h_{\text{biofilm}}\, I_{\text{cell}}}{A_{\text{biofilm}}} \tag{93}$$

$$N_{\text{cells}} = \frac{I_{\text{cell}}\, \xi_{\text{eMtrQ}}\, \xi_{\text{eMtr}}}{e\, \dot{\nu}_e} \tag{94}$$

$$V_{\text{biofilm}} = A_{\text{biofilm}}\, h_{\text{biofilm}} \tag{95}$$

$$V_{\text{biofilm}} = \frac{h_{\text{layer}}\, N_{\text{cells}}\, A_{\text{cell}}}{f_{\text{layer}}} \tag{96}$$

We also considered a monolayer of cells in which the overpotential was set by the direct contact resistance between the electrode and cell. We developed four coupled equations, to solve for the cell current, $I_{\text{cell}}$; the total bias voltage due to the contact resistance $\Delta U_{\text{contact}}$; the total number of cells $N_{\text{cells}}$; and the area of the monolayer of cells $A_{\text{monolayer}}$.

We included the parameters of the individual contact resistance, $R_{\text{contact}}$; the thermodynamic minimum voltage across the cell $U_{\text{redox}}$ set by the half-potentials of Mtr and water splitting at the anode; any additional bias that needs to be applied across the cell $U_{\text{bias}}$; the electron uptake rate per cell $\dot{\nu}_e$; the Faradaic efficiency of transfer of electrons from the electrode to Mtr $\xi_{\text{eMtr}}$; the Faradaic efficiency of transfer of electrons from Mtr to the quinone pool $\xi_{\text{eMtrQ}}$; the fraction of the area of the monolayer of cells occupied by cells, or fill factor $f_{\text{monolayer}}$; and the cross sectional area of a cell $A_{\text{cell}}$.

$$I_{\text{cell}} = \frac{P_{\text{e, avail}}}{U_{\text{redox}} + U_{\text{bias}} + \Delta U_{\text{contact}}} \tag{97}$$

$$\Delta U_{\text{contact}} = \frac{I_{\text{cell}}\, R_{\text{contact}}}{N_{\text{cells}}\, n_{\text{contacts}}} \tag{98}$$

$$N_{\text{cells}} = \frac{I_{\text{cell}}\, \xi_{\text{eMtrQ}}\, \xi_{\text{eMtr}}}{e\, \dot{\nu}_e} \tag{99}$$

$$A_{\text{monolayer}} = N_{\text{cells}}\, A_{\text{cell}} / f_{\text{monolayer}} \tag{100}$$

These coupled equations were solved using the PYTHON SCIPY.OPTIMIZE toolkit.

### 9 Representative Extracellular Matrix Resistivities and Thicknesses

What sorts of resistivities (or its inverse, conductivities) could we expect for a conductive extracellular matrix? We surveyed the literature and found a wide range of values for both biological and non-biological material that could be used to construct a conductive matrix to supply carbon fixing cells with charge.

Malvankar *et al.* measured the resistivity of biofilms produced by four strains of *Geobacter sulfurreducens* and found them to range from 216 to 8,952 $\Omega$ cm with thicknesses between 40 to 130 $\mu$m [31, 32]. Similarly, Yates *et al.* measured the conductivity of *G. sulfurreducens* biofilms to be 200 $\Omega$ cm [33]. The resistivity of *Shewanella oneidensis* by contrast is reported to be 2000 $\Omega$ cm [34]. While *Shewanella oneidensis* is generally considered to produce thinner biofilms than *G. sulfurreducens* [35], Renslow *et al.* were able to produce *S. oneidensis* biofilms that were 350 to 450 $\mu$m thick [36]. As a comparison point, Malvankar *et al.* found the resistivity of *E. coli* biofilms to be $10^5$ $\Omega$ cm [35]. Hung *et al.* measured the thickness of *E. coli* biofilms to be approximately 34 $\mu$m [37].

El-Naggar *et al.* found the resistivity of fixed and dried individual conductive nanowires from *Shewanella oneidensis* to be 1 $\Omega$ cm with lengths up to 600 nm [38].

High conductivity (low resistivity) and high thickness. *Geobacter* [32] and *Shewanella* conductive biofilms are typically several tens of microns thick, the recent discovery of cable bacteria that produce conductive filaments up to 1.5 cm long [39] might revolutionize the potential of conductive biofilms in electromicrobial production. Recently Meysman *et al.* showed that individual conductive filaments from the cable bacteria *Thiofilum flexile* can have resistivities as low as $1.3 \times 10^{-2}$ $\Omega$ cm, over lengths of up to 10.1 mm [40].

Renslow *et al.* predict a linear relationship between maximum biofilm thickness and conductivity [34]. For a resistivity of $10^4 \Omega$ cm, a maximum thickness of 1 cm was predicted, while for a conductivity of $10^{12}$ $\Omega$ cm a maximum thickness of 1 $\mu$m was predicted [34].

As a point of comparison, undoped polyanniline has a resistivity of $3.73\times10^{10}\,\Omega\,\mathrm{cm}$ [41]. Doping with HBr reduces the resistivity to $2.17\times10^{6}\,\Omega\,\mathrm{cm}$ [41]. Polypyrrole, depending upon preparation technique, can have a resistivity as high as $10^{9}\,\Omega\,\mathrm{cm}$ to as low as $1.1\times10^{-2}\,\Omega\,\mathrm{cm}$ [42]. Yu *et al.* constructed an artificial conductive matrix for *S. oneidensis* with a resistivity of $312\,\Omega\,\mathrm{cm}$ with a thickness of 200 to 600 $\mu$m [43].

The results of this section are summarized in **Tables S3** and **S4**.

### 10 Efficiency of Electromicrobial Production with *ex vivo* Electrochemical $CO_2$-fixation

#### 10.1 Definition of Efficiency in Electrochemical $CO_2$-fixation Schemes

For the electrochemical $CO_2$ fixation schemes (**Fig. 1**), we split the available electrical power, $P_{\mathrm{e,\ avail}}$, between (for example the output of a solar PV) between two electrochemical cells, one that electrochemically fixes $CO_2$ and the other that supplies electroactive microbes with electrons to reduce the initial fixation product to fuel. The efficiency of this scheme is the total fuel production rate in the second cell, $\dot{N}_{\mathrm{2f}}$, multiplied by the energy content of each fuel molecule, $E_{\mathrm{fuel}}$, relative to the total input power to the system, $P_{\mathrm{input,\ total}}$,

$$\eta = \frac{\dot{N}_{\mathrm{2f}}\, E_{\mathrm{fuel}}}{P_{\mathrm{input,\ total}}}. \tag{101}$$

The total fuel production rate in the second cell is equal to the current in cell two, $I_2$, multiplied by the Faradaic efficiency of the electrochemical reaction in the second cell, $\xi_{\mathrm{I2}}$, divided by the charge needed to be added in the second cell to synthesize a fuel molecule from the fixation production produced in the first cell, $e\,\nu_{\mathrm{e,\ add}}$,

$$\dot{N}_{\mathrm{2f}} = \frac{I_2\, \xi_{\mathrm{I2}}}{e\,\nu_{\mathrm{e,\ add}}}. \tag{102}$$

By conservation of energy, the total electrical power available for driving electrochemical reactions, $P_{\mathrm{e,\ avail}}$, is equal to the sum of the power in both cells,

$$P_{\mathrm{e,\ avail}} = P_{\mathrm{cell\ 1}} + P_{\mathrm{cell\ 2}}, \tag{103}$$

$$P_{\mathrm{e,\ avail}} = I_1\, \Delta U_{\mathrm{cell\ 1}} + I_2\, \Delta U_{\mathrm{cell\ 2}} \tag{104}$$

The voltage across the first cell, the electrochemical $CO_2$-fixing cell is, the minimum thermodynamic potential difference required across the cell, $U_{\mathrm{cell\ 1,\ 0}}$, and a bias voltage required to drive the reaction,

$$\Delta U_{\mathrm{cell\ 1}} = U_{\mathrm{cell\ 1,\ 0}} + U_{\mathrm{cell\ 1,\ bias}}. \tag{105}$$

The voltage across the first cell, the bio-electrochemical cell is, the minimum thermodynamic potential difference required across the cell, $U_{\mathrm{cell\ 2,\ 0}}$, typically the difference between water splitting and either the $H_2$-reducing potential or the Mtr complex midpoint redox potential, and a bias voltage required to drive the reaction,

$$\Delta U_{\mathrm{cell\ 2}} = U_{\mathrm{cell\ 2,\ 0}} + U_{\mathrm{cell\ 2,\ bias}}. \tag{106}$$

For both $H_2$ and EET supplied microbes the current in the second cell,

$$
\begin{aligned}
I_2 &= \frac{P_{\text{e, avail}}}{\Delta U_{\text{cell 1}}\ (I_1/I_2) + \Delta U_{\text{cell 2}}}, &(107)\\
\dot{N}_{2\text{f}} &= \frac{P_{\text{e, avail}}\,\xi_{\text{I2}}}{e\,\nu_{\text{e, add}}\ (\Delta U_{\text{cell 1}}\ (I_1/I_2) + \Delta U_{\text{cell 2}})}. &(108)
\end{aligned}
$$

Thus, the efficiency of the scheme,

$$
\eta = \frac{P_{\text{e, avail}}\,E_{\text{fuel}}\,\xi_{\text{I2}}}{e\,\nu_{\text{e, add}}\ (\Delta U_{\text{cell 1}}\ (I_1/I_2) + \Delta U_{\text{cell 2}})\ P_{\text{input, total}}}. \tag{109}
$$

#### 10.2 Division of Current Between First and Second Electrochemical Cells

We propose that the ratio of currents between the first and second cell is determined by the ratio of electrons needed to reduce $CO_2$ to the primary electrochemical fixation product in the first cell, and the number of electrons needed to then fully reduce that product to a fuel in the second cell.

As before, for systems where the microbe is supplied by $H_2$ as in **Equation S18** , the number of electrons needed to synthesize a fuel molecule from the lightly reduced electrochemical fixation product of the first cell is,

$$
\begin{aligned}
\nu_{\text{e, add}, \text{H}_2} = {} & 2\,\nu_{\text{f, NADH}} + 2\,\nu_{\text{f, Fd}} \\
& + \nu_{\text{f, ATP}} \frac{\text{ceil}\left(\Delta G_{\text{ATP/ADP}}/e\,\Delta U_{\text{membrane}}\right)}{\text{floor}\left(\left(U_{\text{H}_2} - U_{\text{Acceptor}}\right)/\Delta U_{\text{membrane}}\right)}.
\end{aligned} \tag{110}
$$

While for systems where the microbe is supplied by EET as in **Equation S90**,

$$
\begin{aligned}
\nu_{\text{e, add}, \text{EET}} = {} & 2\,\nu_{\text{f, NADH}} + 2\,\nu_{\text{f, Fd}} \\
& + \nu_{\text{f, ATP}} \frac{\text{ceil}\left(\Delta G_{\text{ATP/ADP}}/e\,\Delta U_{\text{membrane}}\right)}{\text{floor}\left(\left(U_{\text{Q}} - U_{\text{Acceptor}}\right)/\Delta U_{\text{membrane}}\right)} \\
& + 2\,\nu_{\text{f, NADH}} \frac{\text{ceil}\left(U_{\text{NADH}} - U_{\text{Q}}/\Delta U_{\text{membrane}}\right)}{\text{floor}\left(\left(U_{\text{Q}} - U_{\text{Acceptor}}\right)/\Delta U_{\text{membrane}}\right)} \\
& + 2\,\nu_{\text{f, Fd}} \frac{\text{ceil}\left(U_{\text{Fd}} - U_{\text{Q}}/\Delta U_{\text{membrane}}\right)}{\text{floor}\left(\left(U_{\text{Q}} - U_{\text{Acceptor}}\right)/\Delta U_{\text{membrane}}\right)}.
\end{aligned} \tag{111}
$$

However, in both cases the NAD(P)H, Fd and ATP requirements are lower than for fuel synthesis from $CO_2$ as the product of the first cell is already partially reduced.

The current in the first cell is equal to the $CO_2$-fixation rate in the first cell, $\dot{C}_{\text{p}}$, multiplied by the charge needed to synthesize the primary electrochemical $CO_2$-fixation product, $e\,\nu_{\text{ep}}$, divided by the number of carbons in each product, $\nu_{\text{Cp}}$, and the Faradaic efficiency of the first electrochemical reaction, $\xi_{\rm I1}$ (as this is an upper estimate on efficiency, we typically assume that $\xi_{\rm I1} = 1.0$),

$$\dot{N}_{\rm p} = \frac{\dot{C}_{\rm p}}{\nu_{\rm Cp}}, \quad (112)$$

$$I_1 = \frac{\dot{C}_{\rm p}\, e\, \nu_{\rm ep}}{\nu_{\rm Cp}\, \xi_{\rm I1}}, \quad (113)$$

$$I_1 = \frac{\dot{N}_{\rm p}\, e\, \nu_{\rm ep}}{\xi_{\rm I1}} \quad (114)$$

The current flowing to the second cell needs to be adequate to reduce the primary fixation product. Thus,

$$I_2 = \frac{\dot{N}_{\rm p}\, \xi_{\rm C}\, e\, \nu_{\rm e,\ add}}{\nu_{\rm P}\, \xi_{\rm I2}}, \quad (115)$$

$$I_2 = \frac{\dot{C}_{\rm p}\, \xi_{\rm C}\, e\, \nu_{\rm e,\ add}}{\nu_{\rm Cp}\, \nu_{\rm P}\, \xi_{\rm I2}}. \quad (116)$$

Thus, the ratio of currents in the first and second cells,

$$\frac{I_1}{I_2} = \frac{\nu_{\rm P}\, \nu_{\rm ep}\, \nu_{\rm Cp}\, \xi_{\rm I2}}{\xi_{\rm I1}\, \xi_{\rm C}\, \nu_{\rm e,\ add}}. \quad (117)$$

Thus, the efficiency of electrochemical $CO_2$-fixation electromicrobial production schemes is in general,

$$\eta = \frac{P_{\rm e,\ avail}\, E_{\rm fuel}\, \xi_{\rm I2}}{e\, \nu_{\rm e,\ add} \left( \Delta U_{\rm cell\ 1} \left( \frac{\nu_{\rm P}\, \nu_{\rm ep}\, \nu_{\rm Cp}\, \xi_{\rm I2}}{\xi_{\rm I1}\, \xi_{\rm C}\, \nu_{\rm e,\ add}} \right) + \Delta U_{\rm cell\ 2} \right) P_{\rm input,\ total}}. \quad (118)$$

### 11 Carbon, ATP and Reductant demands for Biofuel Synthesis

We estimated the number of NAD(P)H, Ferredoxin (Fd) and ATP required for synthesis of the biofuels isopropanol and butanol for all known naturally-occurring carbon-fixation pathways, the synthetic CETCH [13] pathways, and the synthetic Formolase formate assimilation pathway [44].

In each case, we calculated the number of NAD(P)H, Fd and ATP needed to produce the primary metabolic output of each pathway (pyruvate, glyceraldehyde 3-phosphate (G3P), or acetyl-CoA) either from the primary literature or the BRENDA database [45, 46]. We then calculated the number of NAD(P)H, Fd and ATP required to convert this intermediate to acetyl-CoA, and then to either isopropanol or butanol [47–49]. Full details are included in **Table S2**. All numbers were cross-checked by reference to the BRENDA database [45].

[18] Reference air mass 1.5 spectra.

[19] Jenny Nelson. *The Physics of Solar Cells*. Imperial College Press, 2003.

[20] Joseph H. Noggle. *Physical Chemistry*. Harper Collins, 1996.

[21] Luca Lippi, Lars Bähr, Arne Wüstenberg, Annegret Wilde, and Ralf Steuer. Exploring the potential of high-density cultivation of cyanobacteria for the production of cyanophycin. *Algal Research*, 31:363–366, 2018.

[22] Toshihisa Kanamaru Takeharu Kawano Kenji Tanaka, Ayaaki Ishizaki. Production of poly(D-3-hydroxybutyrate) from $CO_2$, $H_2$, and $O_2$ by high cell density autotrophic cultivation of *Alcaligenes eutrophus*. *Biotechnology and Bioengineering*, 45:268–275, 1995.

[23] Charles Abbas. Personal communication, September 2019.

### Supplementary Information Figures For:

### Constraints on the Efficiency of Electromicrobial Production

*Farshid Salimijazi[a], Jaehwan Kim[b,c], Alexa M. Schmitz[a], Richard Grenville[d], Andrew Bocarsly[b], and Buz Barstow[a,1]*

[a]*Department of Biological and Environmental Engineering, Cornell University, Ithaca, NY 14853, USA;*

[b]*Department of Chemistry, Princeton University, Princeton, NJ 08540, USA;*

[c]*Present Address: Department of Chemistry and Chemical Biology, Cornell University, Ithaca, NY 14853, USA;*

[d]*Philadelphia Mixing Solutions Ltd., Palmyra, PA 17078, USA*

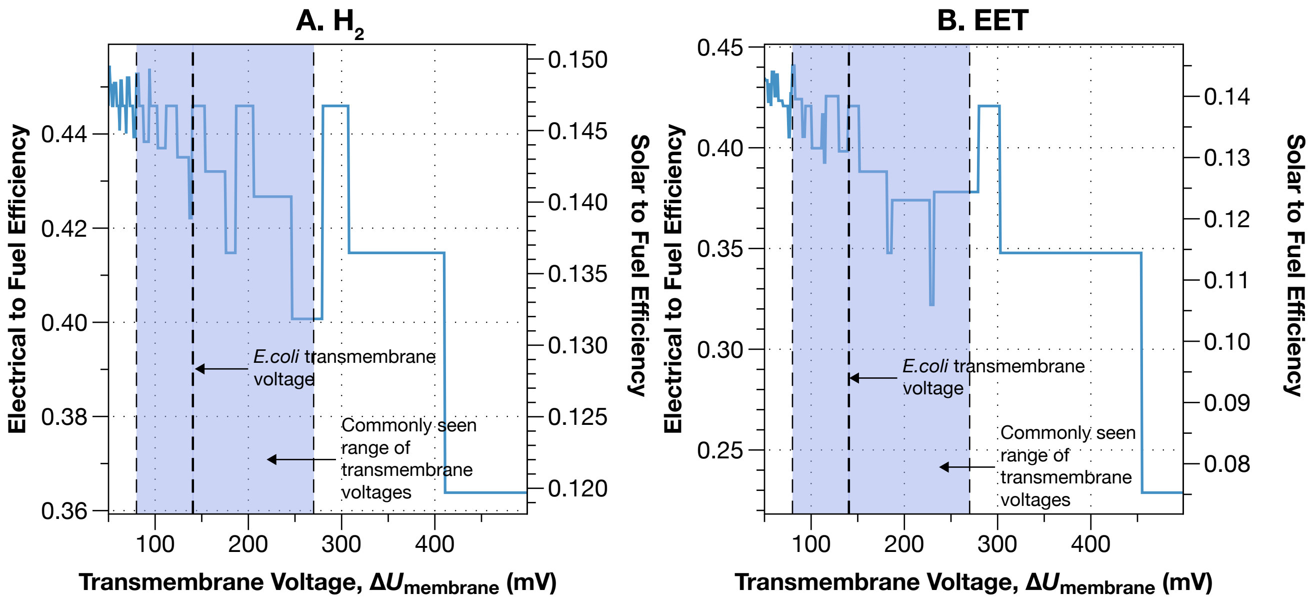


**Fig. S1.** Increasing transmembrane voltage reduces overall efficiency for $H_2$- and EET-mediated electromicrobial production schemes. (**A**) $H_2$-mediated electromicrobial production using the Calvin cycle, and (**B**) EET-mediated electromicrobial production using the Calvin cycle. This plot can be recreated with the sfig-transmembrane.py program and corresponding input file in the rewiredcarbon package.

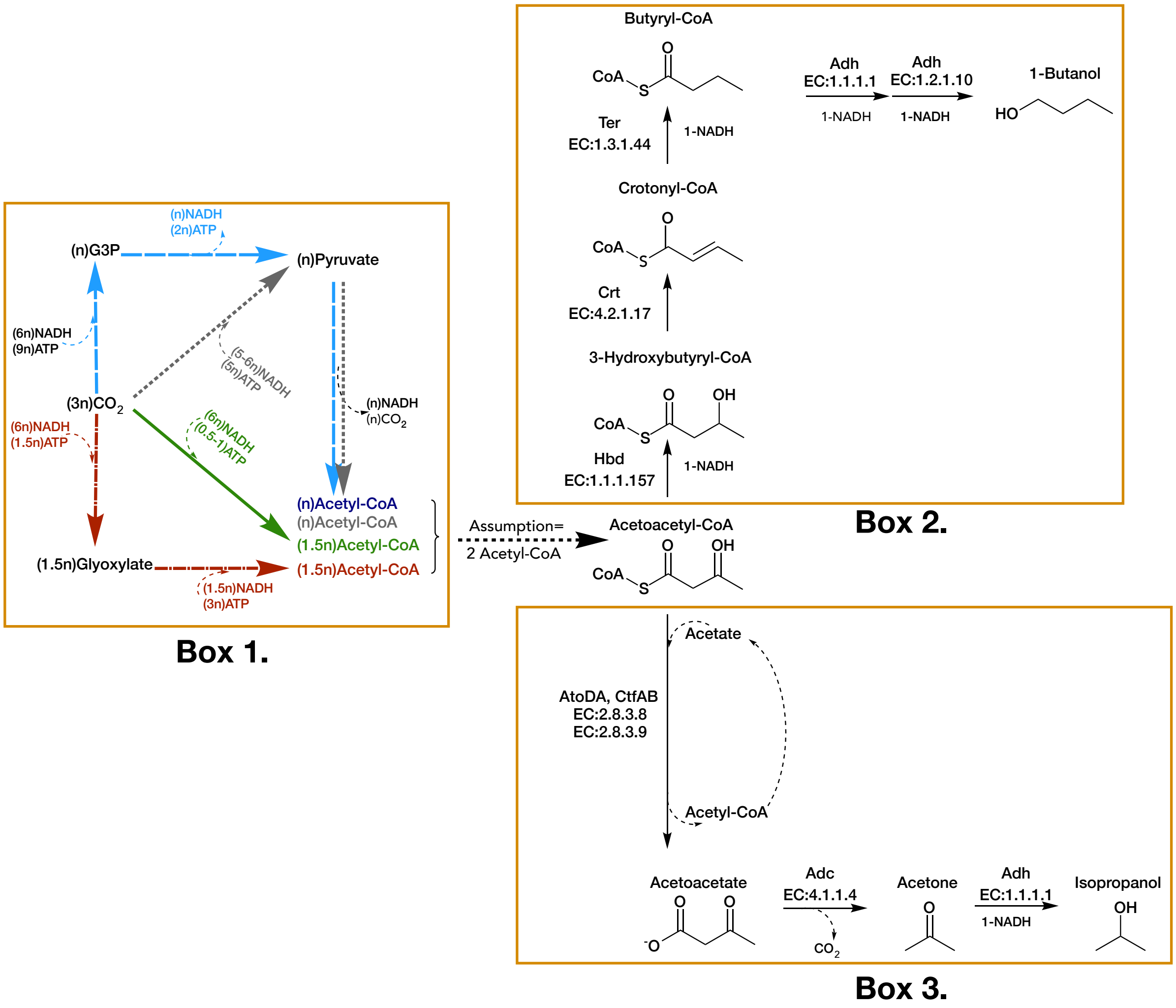


**Fig. S2.** Metabolic pathways for isopropanol and butanol production. Box 1 shows acetyl-CoA production as an intermediate product derived from natural and synthetic $CO_2$ fixation pathways. $CO_2$ is assimilated through; the Calvin cycle (Blue dashed line) first leads to the formation of G3P, which is converted to pyruvate and further to acetyl-CoA; WL and rTCA (Green solid line) produces acetyl-CoA; 3HB and 3HB-4HP (Grey dotted line) leads to the formation of pyruvate and then acetyl-CoA; and CETCH (Red dash-dotted line) leads to the formation of glyoxylate then acetyl-CoA. The ATP and NAD(P)H consumption/production is color-coded for each pathway. Box 2 and 3 show the enzymatic pathway for conversion of two acetyl-CoA molecules to a molecule of 1-butanol or isopropanol, respectively.

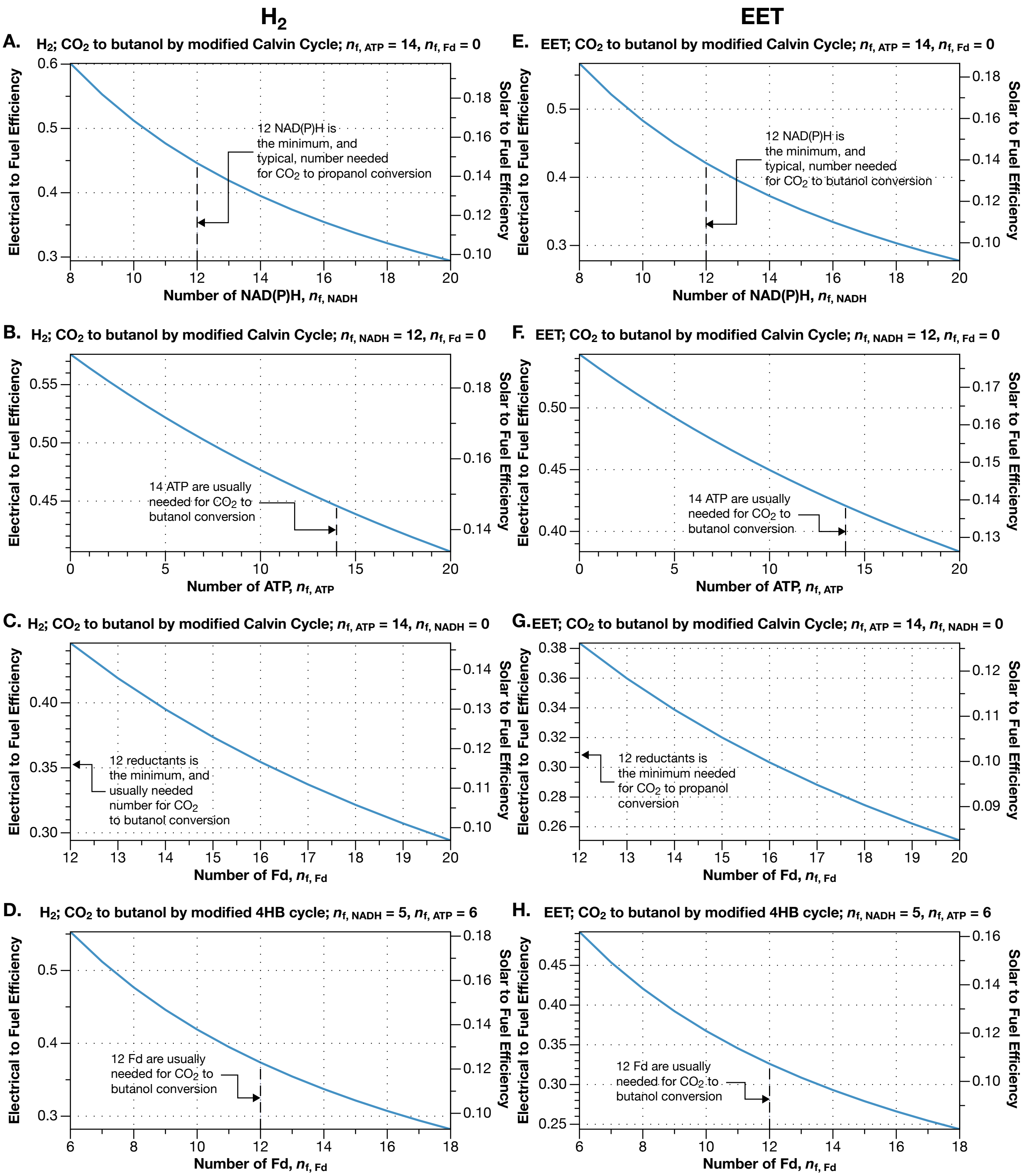


**Fig. S3.** Effect of varying NAD(P)H, Ferredoxin and ATP and requirements for electricity to butanol conversion efficiency. (**A**-**D**) $H_2$-mediated rewired carbon fixation, and (**E**-**H**) EET-mediated rewired carbon fixation. In all cases the transmembrane voltage is 140 mV. This plot can be recreated with the sfig-NADH_ATP_Fd.py program and corresponding input file in the rewiredcarbon package.

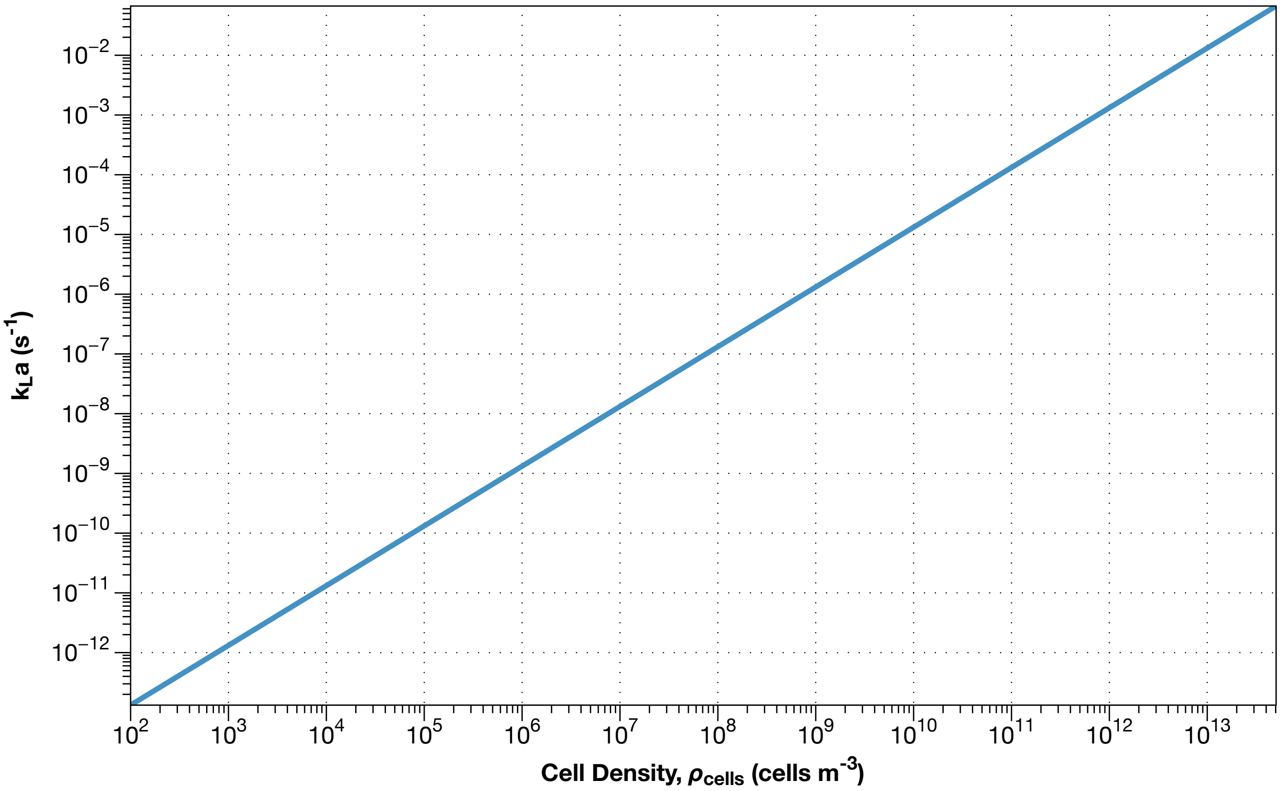


**Fig. S4.** Volumetric transfer rate ($k_L$a) of $H_2$ rises with increasing cell density and decreasing volume. This figure can be recreated with the fig-h2agitation-B-and-C.py program and corresponding input file in the rewiredcarbon package.

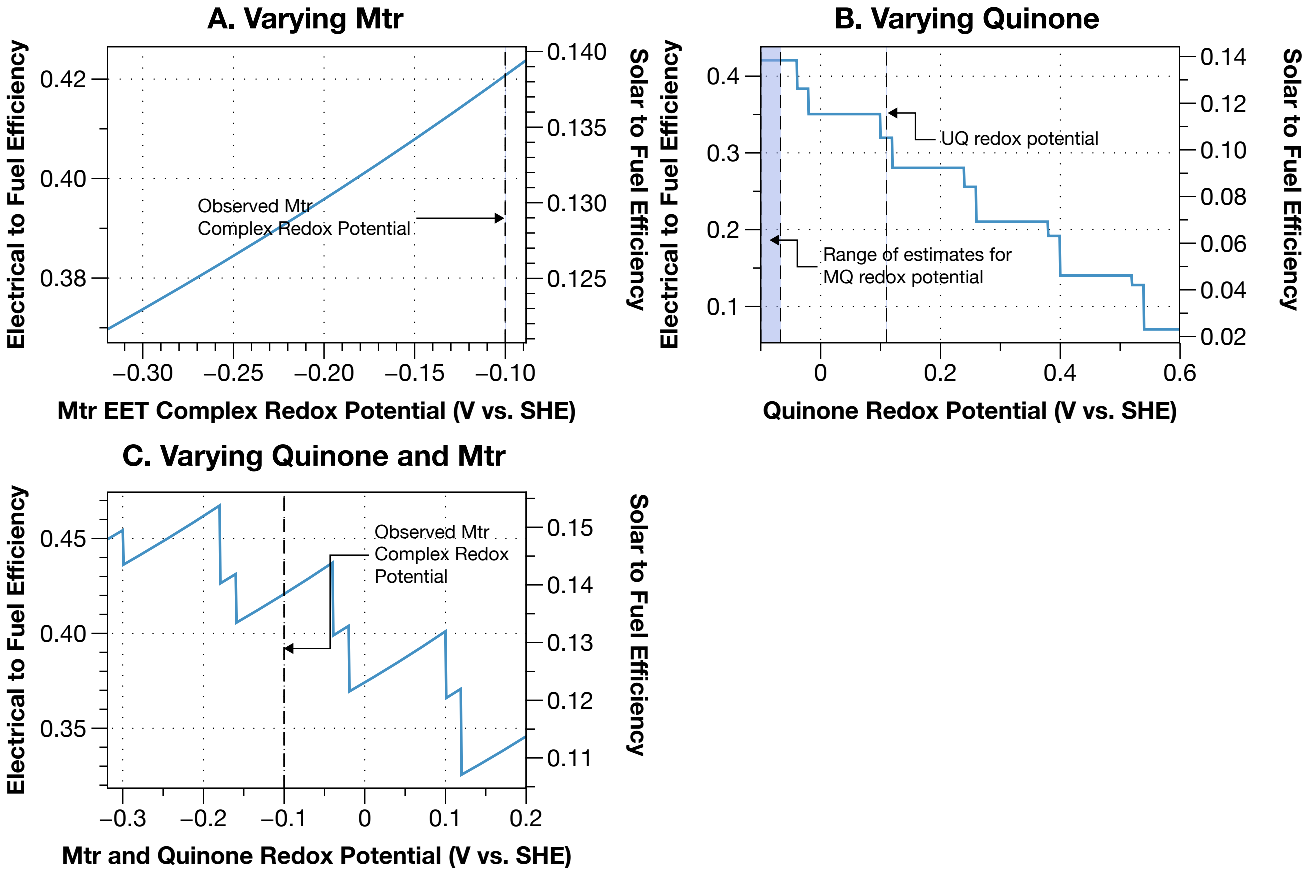


**Fig. S5.** Effect of changing the redox potential of components of the Mtr complex on electrical to fuel efficiency. In each case the anode voltage is set to the water/oxygen couple at 0.82 V with an overpotential of 0.47 V. (**A**) Changing the MtrC redox potential. The redox potential of quinone is fixed at -0.0885 V, while the cathode redox potential is fixed to that of Mtr with constant overpotential of 0.2 V (**B**) Changing the quinone redox potential. The MtrC redox potential is fixed at -0.1 V, while the cathode is set to +0.1 V (a 0.2 V overpotential) (**C**) Effect of simultaneously varying the MtrC and quinone redox potentials on efficiency. The cathode redox potential is fixed to that of Mtr with constant overpotential of 0.2 V. This plot can be recreated with the sfig-vMtr-quinone -A, -B, and -C.py programs and corresponding input files in the rewiredcarbon package.

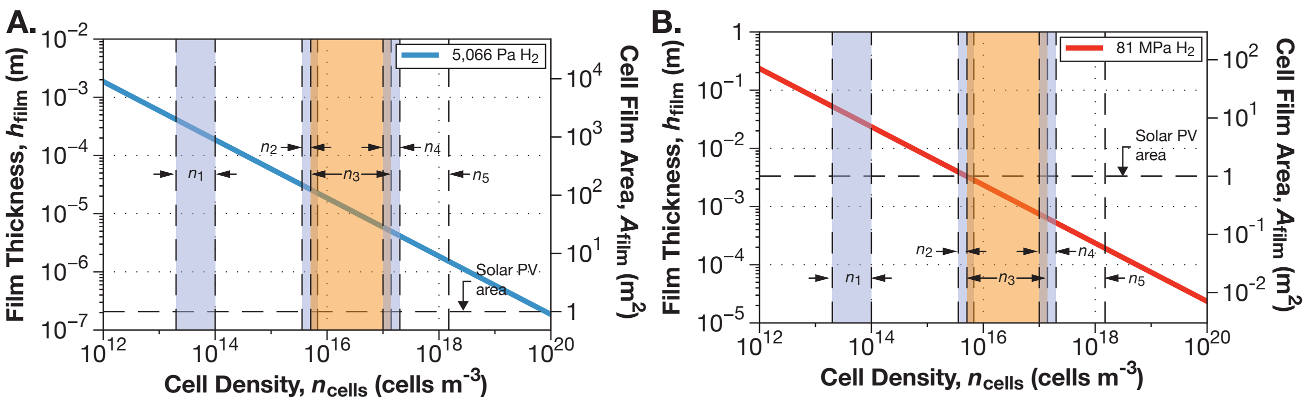


**Fig. S6.** Maximum height of cell culture that can be supplied with $H_2$ by diffusion and corresponding area of culture needed to convert 330 W of electrical power to butanol. In this figure, we have separated out the high and low pressure film height and area curves in **Fig. 4B** in the main text into separate subplots. This allows us to represent the area and height of the thin-film culture with a single curve and a set of inversely related axes. (**A**) Maximum height and minimum area of cell culture that can be supplied with by diffusion from headspace with at $H_2$ partial pressures of 5,066 Pa (5% of atmospheric pressure). (**B**) Maximum height and minimum area that can be supplied by diffusion with $H_2$ at a partial pressure of 81 MPa (80% of 1,000 atmospheres). Five important cell density regimes are noted in both panels: $n_1$: laboratory grown cultures of *E. coli* in exponential phase; $n_2$: cyanobacteria grown to maximum density; $n_3$: cultures of *E. coli* at saturating density; $n_4$: $H_2$-oxidizing microbes grown to maximum density; and $n_5$: and saturating cultures of industrially-grown yeast (**SI Text 5**).

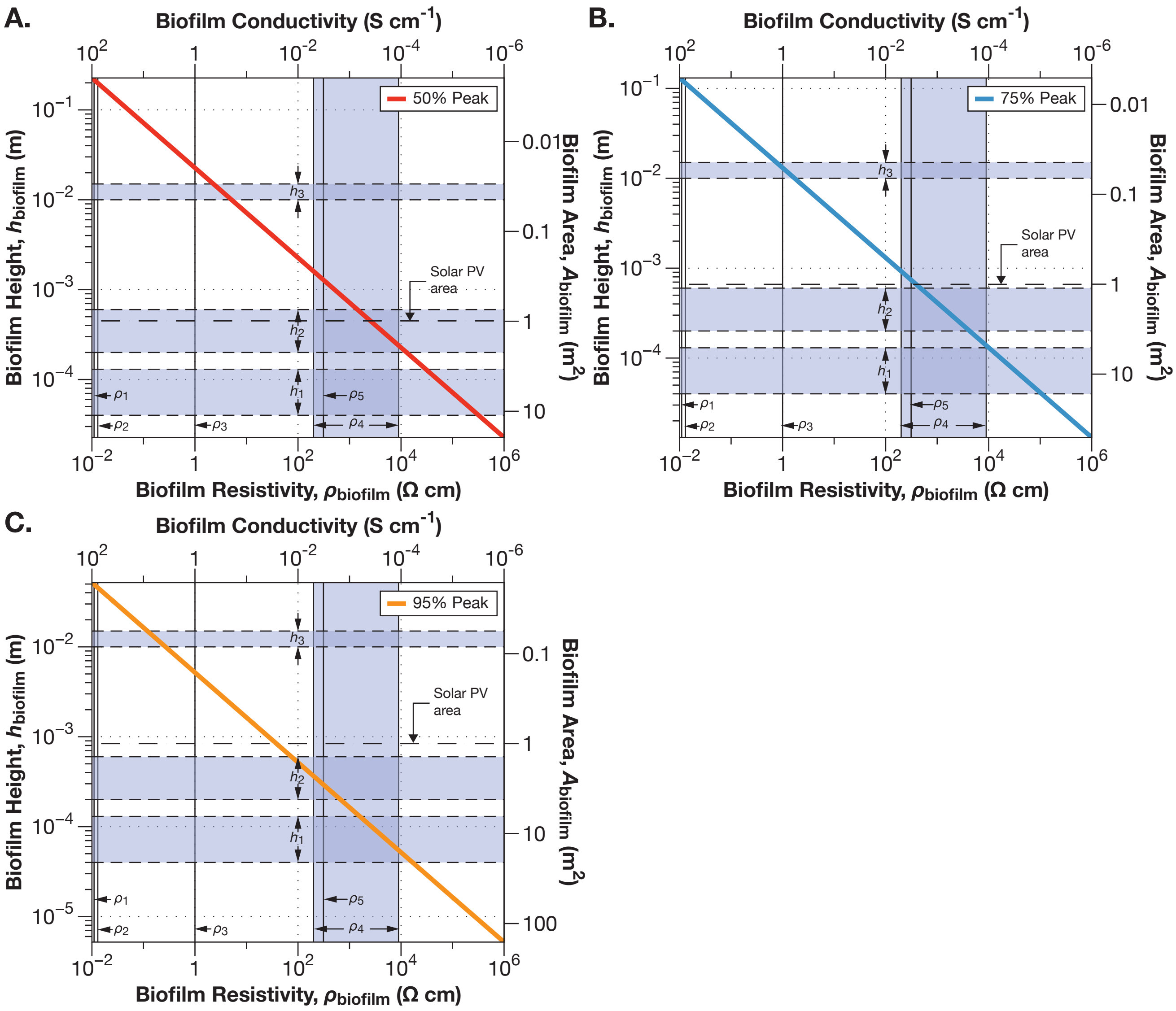


**Fig. S7.** Maximum biofilm thickness and minimum area needed to achieve 50, 75 and 95% of peak efficiency. In this figure, we have separated out the conductive extracellular matrix (ECM) thickness and area curves in **Fig. 6B** in the main text into separate subplots for each efficiency. This allows us to represent the area and height of the ECM with a single curve and a set of inversely related axes. ECM thickness and area for (**A**) 50% of peak efficiency; (**B**) 75% of peak efficiency; and (**C**) 95% of peak efficiency. Representative conductive matrix resistivities and heights are marked in each subplot: $\varrho_1$: high conductivity polypyrrole; $\varrho_2$: individual cable bacteria filaments; $\varrho_3$: individual *S. oneidensis* nanowires; $\varrho_4$: bulk *G. sulfurreducens* and *S. oneidensis* biofilm resistivities; $\varrho_5$: polypyrrole conductive matrix for *S. oneidensis*; $\varrho_6$: bulk *E. coli* biofilm; $\varrho_7$: HBr doped polyaniline; $\varrho_8$: low conductivity polypyrrole; $h_1$: *G. sulfurreducens* biofilms; $h_2$: polypyrrole conductive matrix for *S. oneidensis*; $h_3$: cable bacteria biofilms and individual filaments (**SI Text 9** and **Tables S3** and **S4**).

### Supplementary Table 1: Symbols Used in this Article

| Symbol | Definition | Unit | Macro |
|---|---|---|---|
| **Physical Constants** | | | |
| $N_\mathrm{A}$ | Avogadro constant | $\mathrm{mol}^{-1}$ | `\avogadroConstant` |
| $e$ | Elementary charge | A s | `\elementaryCharge` |
| $R$ | Ideal gas constant | $\mathrm{J\,K\,mol}^{-1}$ | `\idealGasConstant` |
| $F$ | Faraday constant | $\mathrm{A\ s\ mol}^{-1}$ | `\faradayConstant` |
| $k_\mathrm{B}$ | Boltzmann constant | $\mathrm{J\,K}^{-1}$ | `\boltzmannConstant` |
| **Widely Used Variables** | | | |
| $V_\mathrm{cells}$ | Cell culture volume | $\mathrm{m}^3$ | `\cellCultureVolume` |
| $N_\mathrm{cells}$ | Total number of cells | # | `\totalCells` |
| $E_\mathrm{fuel}$ | Energy content per fuel molecule | J | `\energyPerFuelMolecule` |
| $\mathrm{n_{cells}}$ | Cell culture density | $\mathrm{m}^{-3}$ | `\cellDensity` |
| $\Delta U_\mathrm{cell}$ | Whole electrochemical cell voltage | V | `\voltageCell` |
| $I_\mathrm{cell}$ | Whole electrochemical cell current | A | `\currentCell` |
| $I_\mathrm{cell,\ bio}$ | Whole electrochemical cell current available to biological processes | A | `\currentCellBio` |
| $\dot{N}_\mathrm{fuel}$ | Total fuel molecule generation rate in electrochemical cell | $\mathrm{s}^{-1}$ | `\totalFuelProductionRate` |
| $T$ | Temperature | K | `\temperature` |
| $l_\mathrm{cell}$ | Length of cell | m | `\cellLength` |
| $d_\mathrm{cell}$ | Diameter of cell | m | `\cellDiameter` |
| $A_\mathrm{cell}$ | Cross-sectional area of cell | $\mathrm{m}^2$ | `\crossSectionalAreaCell` |
| $\eta$ | Overpotential | V | `\overpotential` |
| $U_\mathrm{redox}$ | Electrochemical cell redox reaction voltage | V | `\voltageRedox` |
| $U_\mathrm{bias}$ | Electrochemical cell additional bias | V | `\voltageBias` |
| $U_{\mathrm{H_2}}$ | Electrochemical cell proton to $H_2$ reduction reaction | V | `\voltageHydrogen` |
| $U_{\mathrm{O_2}}$ | Electrochemical cell water splitting reaction | V | `\voltageOxygen` |
| $U_{\mathrm{H_2O/O_2}}$ | Electrochemical cell water splitting reaction | V | `\voltageWaterOxygen` |
| $P_\mathrm{input,\ total}$ | Integrated total power input to system (usually total solar power) | W | `\totalInputPower` |
| $P_\mathrm{e,\ total}$ | Total electrical power available to electrochemical cell(s) | W | `\totalElectricalPower` |
| $P_\mathrm{cell\ 1}$ | Electrical power directed to electrochemical cell 1 | W | `\powerCellOne` |
| $P_\mathrm{cell\ 2}$ | Electrical power directed to electrochemical cell 2 | W | `\powerCellTwo` |
| $P_\mathrm{e,\ total}$ | Total electrical power available to electrochemical cell(s) | W | `\totalElectricalPower` |
| $P_\gamma$ | Integrated solar power density | $\mathrm{W\,m}^{-2}$ | `\solarPower` |
| $\eta$ | General overall efficiency | # | `\generalEfficiency` |
| **Reductant and Energy Carrier Demand for Fuel or Energy Storage Molecule Synthesis Variables** | | | |
| $\dot{\nu}_e$ | Electron demand rate per cell | $\mathrm{s}^{-1}$ | `\electronRatePerCell` |

| | | | |
|---|---|---|---|
| $\nu_{\mathrm{ef,\,NADH}}$ | Number of electrons needed to reduce the NAD(P)H necessary to synthesize one fuel molecule from $CO_2$ | # | \electronsNADHforfuel |
| $\nu_{\mathrm{ef,\,ATP,\,EET}}$ | Number of electrons needed to be sent downhill in energy to synthesize the ATP to make one molecule of fuel from $CO_2$ | # | \electronsDownATPforFuelInEET |
| $\nu_{\mathrm{ef,\,NADH,\,down}}$ | Number of electrons needed to be sent downhill to raise $\nu_{\mathrm{ef,\,NADH}}$ electrons in energy | # | \electronsDownNADHforFuel |
| $\nu_{\mathrm{ef,\,Fd,\,down}}$ | Number of electrons needed to be sent downhill to raise $\nu_{\mathrm{ef,\,Fd}}$ electrons in energy | # | \electronsDownFdForFuel |
| $\nu_{\mathrm{ef,\,ATP,\,H_2}}$ | Number of electrons needed to be sent downhill to synthesize the ATP to make one molecule of fuel from $CO_2$ through $H_2$ | # | \electronsDownATPforFuelinH |
| $\nu_{\mathrm{ef}}$ | Total number of electrons needed to synthesize one molecule of fuel from $CO_2$ | # | \electronsPerFuel |
| $\nu_{\mathrm{ef,\,H_2}}$ | Total number of electrons needed to make one fuel molecule in a $H_2$-mediated system | # | \electronsPerFuelinH |
| $\nu_{\mathrm{ef,\,EET}}$ | Total number of electrons needed to make one fuel molecule in an EET-mediated system | # | \electronsPerFuelInEET |
| $\nu_{\mathrm{ep}}$ | Number of electrons needed per carbon to reduce $CO_2$ to primary fixation product in cell 1 | # | \electronsPerPrimaryFix |
| $\nu_{\mathrm{e,\,add,\,H_2}}$ | Number of electrons per carbon to add to primary fixation production to reduce to biofuel in $H_2$ | # | \electronsToAddinH |
| $\nu_{\mathrm{e,\,add,\,EET}}$ | Number of electrons per carbon to add to primary fixation production to reduce to biofuel in EET | # | \electronsToAddinEET |
| $\nu_{\mathrm{e,\,add}}$ | Number of electrons per carbon to add to primary fixation production to reduce to biofuel | # | \electronsToAdd |
| $\nu_{\mathrm{f,\,Fd}}$ | Number of Ferredoxin molecules needed to synthesize one energy storage molecule from either $CO_2$ or an electrochemical $CO_2$ fixation product | # | \FdForFuel |
| $\nu_{\mathrm{f,\,NADH}}$ | Number of NAD(P)H molecules needed to synthesize one energy storage molecule from either $CO_2$ or an electrochemical $CO_2$ fixation product | # | \NADHForFuel |
| $\nu_{\mathrm{f,\,ATP}}$ | Number of ATP molecules needed to synthesize one energy storage molecule from either $CO_2$ or an electrochemical $CO_2$ fixation product | # | \ATPforFuel |
| **Carbon Demand for Fuel or Energy Storage Molecule Synthesis Variables** | | | |
| $\nu_{\mathrm{Cf}}$ | Number of carbon atoms per fuel molecule | # | \carbonsPerFuel |
| $\nu_{\mathrm{Cf,\,fix}}$ | Number of carbon atoms fixed per fuel molecule. $\nu_{\mathrm{Cf,\,fix}} \leq \nu_{\mathrm{Cf}}$ | # | \carbonsFixedPerFuel |

| | | | |
|---|---|---|---|
| $\nu_{\mathrm{P}}$ | Number of electrochemical (primary) carbon-fixation products needed to synthesize one molecule of fuel | # | `\primaryFixPerFuel` |
| $\nu_{\mathrm{Cp}}$ | Number of carbon atoms per primary fixation | # | `\carbonsPerPrimaryFix` |
| $\dot{C}_{\mathrm{p}}$ | Carbon fixation rate by first cell | $s^{-1}$ | `\carbonFixationRateCellOne` |
| $\nu_{\mathrm{fix}}$ | Number of carboxylating enzyme (typically RuBisCO) catalytic units per cell | # | `\cotwoFixEnzymePerCell` |
| $r_{\mathrm{fix}}$ | Turnover rate per catalytic site of primary carboxylating enzyme (typically RuBisCO) | $s^{-1}$ | `\cotwoFixEnzymeRate` |
| $\nu_{\mathrm{efix}}$ | Number of rate-limiting enzymes per cell in pathway that transform electrochemical $CO_2$ fixation product | # | `\primaryFixEnzymesPerCell` |
| $r_{\mathrm{efix}}$ | Turnover rate of rate-limiting enzyme in pathway that transform electrochemical $CO_2$ fixation product | $s^{-1}$ | `\primaryFixEnzymeRate` |
| **Hydrogen-Mediated Rewired Carbon Fixation** | | | |
| $\dot{\nu}_{\mathrm{H_2}}$ | $H_2$ demand rate per cell | $s^{-1}$ | `\hydrogenRatePerCell` |
| $C_{\mathrm{H_2}}$ | $H_2$ concentration in liquid | $m^{-3}$ | `\concentrationHydrogen` |
| $C_{\mathrm{H_2,\ max}}$ | Maximum $H_2$ concentration in liquid | $m^{-3}$ | `\concentrationHydrogenMax` |
| $P_{\mathrm{H_2}}$ | Pressure of $H_2$ in headspace of $H_2$-mediated electrosynthesis reactor | Pa | `\pressureHydrogen` |
| $k_{\mathrm{H_2}}$ | Solubility coefficient of $H_2$ in liquid (water in this case) | $\mathrm{Pa\,m^3\,mol^{-1}}$ | `\solubilityHydrogen` |
| **Hydrogen-Transport by Diffusion** | | | |
| $D_{\mathrm{H_2}}$ | Diffusion coefficient of $H_2$ | $\mathrm{m^2\,s^{-1}}$ | `\diffusionHydrogen` |
| $h_{\mathrm{film}}$ | Height of thin-film culture in electrosynthesis reactor | m | `\hydFilmThickness` |
| $A_{\mathrm{film}}$ | Area of thin-film culture in electrosynthesis reactor supplied by diffusion | $m^2$ | `\hydFilmArea` |
| **Hydrogen-Transport by Agitation** | | | |
| $d_a$ | Agitator diameter | $\mathrm{m\,s^{-1}}$ | `\agitatorDiameter` |
| $u_{\mathrm{SG}}$ | Superficial gas (typically $H_2$) velocity. The volume of gas passing through each square meter of the top of the liquid. | $\mathrm{m\,s^{-1}}$ | `\superficialGasVelocity` |
| $\dot{v}_{\mathrm{gas}}$ | Volumetric transfer rate of gas (typically $H_2$) | $\mathrm{m^3\,s^{-1}}$ | `\volumeTransferRateGas` |
| $J_{\mathrm{gas}}$ | Mass transfer rate of gas into solution (typically $H_2$) | $\mathrm{mol\,s^{-1}}$ | `\massTransferRateGas` |
| $k_{\mathrm{L}}a$ | Mass transfer rate of gas into solution (typically $H_2$) | $s^{-1}$ | `\kLaGas` |
| $A$ | Empirical coefficient used in calculation of $k_{\mathrm{L}}a$. 0.026 for mixing into water, 0.002 for strong ionic solutions. | # | `\Aconstant` |
| $B$ | Empirical coefficient used in calculation of $k_{\mathrm{L}}a$. 0.4 for mixing into water, 0.7 for strong ionic solutions. | # | `\Bconstant` |
| $C$ | Empirical coefficient used in calculation of $k_{\mathrm{L}}a$. 0.5 for mixing into water, 0.2 for strong ionic solutions. | # | `\Cconstant` |
| $\epsilon_{\mathrm{gas}}$ | Empirical coefficient used in calculation of $k_{\mathrm{L}}a$. 0.5 for mixing into water, 0.2 for strong ionic solutions. | $\mathrm{W\,m^{-3}}$ | `\stirPowerDensity` |

| | | | |
|---|---|---|---|
| $\Delta C_{\mathrm{gas}}$ | Difference between peak concentration of gas in equilibrium with bubble and bulk liquid | $\mathrm{mol\,m^{-3}}$ | \gasConcentrationDifference |
| $C_{\mathrm{gas,\ interface}}$ | Concentration of gas in equilibrium with gas bubble | $\mathrm{mol\,m^{-3}}$ | \gasBubbleEquilibrium Concentration |
| $C_{\mathrm{gas,\ bulk}}$ | Concentration of gas (typically $H_2$) in bulk liquid | $\mathrm{mol\,m^{-3}}$ | \gasBulkConcentration |
| $r_{\mathrm{Dh}}$ | Tank diameter to height ratio | # | \tankDiameterToHeightRatio |
| $A_{\mathrm{tank}}$ | Area of tank through which gas passes (typically $H_2$) | $\mathrm{m^2}$ | \tankCrossSectionalArea |
| $h_{\mathrm{tank}}$ | Tank liquid height | m | \tankHeight |
| $d_{\mathrm{tank}}$ | Tank diameter | m | \tankDiameter |
| $P_{\mathrm{e,\ stir}}$ | Electrical power used to stir fluid in electrochemical cell(s) (typically the bio-cell) | W | \stirPower |
| **Extracellular Electron Transfer Conductive Matrix Variables** | | | |
| $h_{\mathrm{biofilm}}$ | Exact thickness of biofilm | m | \heightBiofilm |
| $f_{\mathrm{layer}}$ | Fill factor of layer of biofilm (fraction of layer occupied by cells) | # | \biofilmLayerFillFactor |
| $h_{\mathrm{layer}}$ | Height of layer of biofilm | m | \biofilmLayerHeight |
| $\rho_{\mathrm{biofilm}}$ | Resistivity of biofilm | $\Omega\,\mathrm{m}$ | \resistivityBiofilm |
| $A_{\mathrm{layer}}$ | Surface area of one layer | $\mathrm{m^2}$ | \layerArea |
| $n_{\mathrm{layers}}$ | Number of cell layers in biofilm | # | \numberCellLayers |
| $\Delta U_{\mathrm{biofilm}}$ | Biofilm overpotential | V | \biofilmOverpotential |
| $A_{\mathrm{biofilm}}$ | Area of biofilm | $\mathrm{m^2}$ | \areaBiofilm |
| $V_{\mathrm{biofilm}}$ | Volume of biofilm | $\mathrm{m^3}$ | \biofilmVolume |
| **Extracellular Electron Transfer Conductive Matrix Variables** | | | |
| $n_{\mathrm{contacts}}$ | Number of contact points per cell when making direct contact with electrode | # | \contactsPerCell |
| $R_{\mathrm{contact}}$ | Contact resistance of each direct contact point of the cell with a cathode | $\Omega$ | \contactResistance |
| $f_{\mathrm{monolayer}}$ | Fill factor of monolayer cells making direct contact with cathode | # | \monoLayerFillFactor |
| $\Delta U_{\mathrm{contact}}$ | Contact resistance overpotential | V | \contactResistanceOverpotential |
| $A_{\mathrm{monolayer}}$ | Area of monolayer of cells | $\mathrm{m^2}$ | \areaMonolayer |
| **Intracellular Electrochemical Variables** | | | |
| $U_{\mathrm{Mtr}}$ | Electrochemical cell Mtr complex reduction reaction | V | \voltageMtr |
| $U_{\mathrm{Acceptor}}$ | Redox potential of terminal electron acceptor | V | \voltageAcceptor |
| $U_{\mathrm{NADH}}$ | Redox potential of NADH | V | \voltageNADH |
| $U_{\mathrm{Fd}}$ | Redox potential of Ferredoxin | V | \voltageFd |
| $U_{\mathrm{Q}}$ | Redox potential of quinone molecule | V | \voltageQuinone |
| $\Delta U_{\mathrm{membrane}}$ | Redox potential difference across inner membrane | V | \voltageMembrane |
| **Variables Describing Electrochemical Cells** | | | |
| $I_{\mathrm{H_2}}$ | Hydrogen current in electrochemical cell | A | \hydrogenCurrent |
| $I_{\mathrm{Mtr}}$ | Mtr current in electrochemical cell | A | \currentMtr |
| $I_{\mathrm{Q}}$ | Quinone current in electrochemical cell | A | \currentQuinone |
| $I_{\mathrm{NADH}}$ | Current through NADH | A | \NADHCurrent |
| $I_{\mathrm{Fd}}$ | Current through Ferredoxin | A | \FdCurrent |
| $\xi_{\mathrm{eH_2}}$ | Electrons to hydrogen conversion efficiency | # | \efficiencyElectronsToHyd |
| $\xi_{\mathrm{eMtr}}$ | Current to Mtr conversion efficiency | # | \efficiencyElectronsToMtr |

| | | | |
|---|---|---|---|
| $\xi_{\text{eMtrQ}}$ | Electrons on Mtr to Quinone conversion efficiency | # | \efficiencyElectronsMtrToQuinone |
| $U_{\text{cell 2, cathode, 0}}$ | Thermodynamic minimum potential at the cell two (bio-electrochemical cell) cathode | V | \voltageCellTwoCathode |
| $U_{\text{cell 2, cathode, bias}}$ | Bias voltage at the cell two (bio-electrochemical cell) cathode | V | \voltageCellTwoCathodeBias |
| $U_{\text{cell 2, anode, 0}}$ | Thermodynamic minimum potential at the cell two (bio-electrochemical cell) anode | V | \voltageCellTwoAnode |
| $U_{\text{cell 2, anode, bias}}$ | Bias voltage at the cell two (bio-electrochemical cell) anode | V | \voltageCellTwoAnodeBias |
| $U_{\text{cell 2, bias}}$ | Total bias voltage across cell two (bio-electrochemical cell) | V | \voltageCellTwoBias |
| $\Delta U_{\text{cell 2}}$ | Total voltage across cell two (bio-electrochemical cell) | V | \voltageCellTwo |
| $I_2$ | Current in secondary fuel reduction electrochemical cell | A | \currentCellTwo |
| $U_{\text{cell 1, cathode, 0}}$ | Thermodynamic minimum potential at the cell one (electrochemical $CO_2$ fixation cell) cathode | V | \voltageCellOneCathode |
| $U_{\text{cell 1, cathode, bias}}$ | Bias voltage at the cell one (electrochemical $CO_2$ fixation cell) cathode | V | \voltageCellOneCathodeBias |
| $U_{\text{cell 1, anode, 0}}$ | Thermodynamic minimum potential at the cell one (electrochemical $CO_2$ fixation cell) anode | V | \voltageCellOneAnode |
| $U_{\text{cell 1, anode, bias}}$ | Bias voltage at the cell one (electrochemical $CO_2$ fixation cell) anode | V | \voltageCellOneAnodeBias |
| $\Delta U_{\text{cell 1}}$ | Total voltage across cell one (electrochemical $CO_2$ fixation cell) | V | \voltageCellOne |
| $I_1$ | Current in primary carbon fixation electrochemical cell | A | \currentCellOne |
| $P_{\text{e, avail}}$ | Available electrical power to dispatch to electrochemical cell(s) | W | \availableElectricalPower |
| $\xi_{\text{I1}}$ | Current from solar to first Cell conversion efficiency | # | \efficiencyCurrentToFirstCell |
| $\xi_{\text{I2}}$ | Current from solar to second Cell conversion efficiency | # | \efficiencyCurrentToSecondCell |
| $\xi_{\text{C}}$ | Fixed carbon from first to second cell conversion efficiency | # | \efficiencyCarbonToSecondCell |
| **Solar PV** | | | |
| $U_\gamma$ | Voltage from photovoltaic | V | \photoVoltage |
| $I_\gamma$ | Current from photovoltaic | A | \photoCurrent |
| $G_{\text{SC}}$ | Solar constant | $\text{W m}^{-2}$ | \solarConstant |
| $A_{\text{PV}}$ | Area of solar photovoltaic | $\text{m}^2$ | \areaSolarPV |
| **Efficiency** | | | |
| $\eta_{\text{EF}}$ | Electrical to Fuel conversion efficiency | # | \ElectricalToFuelEfficiency |
| $\eta_{\text{EH}_2\text{F}}$ | Electrical to $H_2$ to Fuel conversion efficiency | # | \ElectricalToHydToFuelEfficiency |
| $\eta_{\text{SE}}$ | Solar to Electrical conversion efficiency | # | \SolarToElectricalEfficiency |
| $\eta_{\text{SH}_2\text{F}}$ | Solar to $H_2$ to Fuel conversion efficiency | # | \SolarToHydToFuelEfficiency |
| $\eta_{\text{EEF}}$ | Electrical to EET to Fuel conversion efficiency | # | \ElectricalToEETtoFuelEfficiency |
| **Uphill Pathway Variables** | | | |

| | | | |
|---|---|---|---|
| $p_{\mathrm{out,\,EET}}$ | Number of protons pumped out by sending one electron downhill to acceptor | # | \numberOfProtonsPumpedOutPer ElectronDownInEET |
| $p_{\mathrm{in,\,NADH}}$ | Number of protons pumped in to send one electron uphill to NAD(P)H | # | \numberOfProtonsPumpedInPer ElectronForNADH |
| $p_{\mathrm{in,\,Fd}}$ | Number of protons pumped in to send one electron uphill to Ferredoxin (Fd) | # | \numberOfProtonsPumpedInPer ElectronForFd |
| $p_{\mathrm{in,\,ATP}}$ | Number of protons pumped in to catalyze ADP to ATP | # | \numberOfProtonsPumpedInForATP |
| $\Delta G_{\mathrm{ATP/ADP}}$ | Gibbs free energy of ATP | $\mathrm{J\,mol^{-1}}$ | \ATPGibbsFreeEnergy |
| $p_{\mathrm{out,\,H_2}}$ | Numbers of protons pumped out by sending one electron downhill to acceptor from $H_2$ | # | \numberOfProtonsPumpedOutPer ElectronDownInH |
| $I_{\mathrm{downhill,\,ATP}}$ | Current sent downhill in energy to terminal electron acceptor to regenerate ATP | A | \downhillATPCurrent |
| $I_{\mathrm{downhill,\,NADH}}$ | Current sent downhill in energy to terminal electron acceptor to produce proton gradient needed to reduce NAD(P)H | A | \downhillNADHCurrent |
| $I_{\mathrm{downhill,\,Fd}}$ | Current sent downhill in energy to terminal electron acceptor to produce proton gradient needed to reduce Ferredoxin (Fd) | A | \downhillFdCurrent |
| $I_{\mathrm{NADH}}$ | Current used to directly reduce NAD(P)H | A | \NADHCurrent |
| $I_{\mathrm{Fd}}$ | Current used to directly reduce Ferredoxin (Fd) | A | \FdCurrent |

**Table S2.** Carbon, ATP and reductant demands for biofuel synthesis for naturally-occurring and synthetic carbon fixation pathways. Detailed discussion is included in **SI Text 11**. Reference numbers refer to main text references. Fd: reduced ferredoxin; G3P: Glyceraldehyde 3-phosphate; 3-PG: 3-Phosphoglyceric acid. NADH is used to denote NADH or NAD(P)H.

| **Pathway** | **Reaction** | **Reference(s)** |
|---|---|---|
| ***Wood-Ljungdahl (WL) Pathway*** | | |
| $CO_2$ to metabolic intermediate | 4 $CO_2$ + 2 ATP + 8 NADH → 2 acetyl-CoA | Berg *et al*. [97] |
| Intermediate to butanol | 2 acetyl-CoA + 4 NADH → 1 butanol | Schadeweg *et al*. [98] |
| *Overall $CO_2$ to butanol* | 4 $CO_2$ + 2 ATP + 12 NADH → 1 butanol | |
| ***Calvin-Benson-Bassham (CBB) Cycle*** | | |
| $CO_2$ to metabolic intermediate | 6 $CO_2$ + 18 ATP + 12 NADH → 2 G3P | Berg *et al*. [97] |
| | 2 G3P → 2 pyruvate + 4 ATP + 2 NADH | Berg *et al*. [97], Schomburg *et al*. [94] |
| | 2 pyruvate → 2 acetyl-coA + 2 NADH + 2 $CO_2$ | Berg *et al*. [97], Schomburg *et al*. [94] |
| Intermediate to propanol | | |
| Intermediate to butanol | 2 acetyl-CoA + 4 NADH → 1 butanol | Schadeweg *et al*. [98] |
| *Overall $CO_2$ to propanol* | 4 $CO_2$ + 14 ATP + 9 NADH → 1 isopropanol + 1 $CO_2$ | |
| *Overall $CO_2$ to butanol* | 6 $CO_2$ + 14 ATP + 12 NADH → 1 butanol + 2 $CO_2$ | |
| ***(CoA)/ethylmalonyl-CoA/hydroxybutyryl-CoA (CETCH) Pathway*** | | |
| $CO_2$ to metabolic intermediate | 4 $CO_2$ + 2 ATP + 8 NADH → 2 glyoxylate | Schwander *et al*. [35] |
| | 2 glyoxylate + 4 ATP + 2 NADH → 2 acetyl-CoA | Yu *et al*. [96] |
| Intermediate to butanol | 2 acetyl-CoA + 4 NADH → 1 butanol | Schadeweg *et al*. [98] |
| *Overall $CO_2$ to butanol* | 4 $CO_2$ + 6 ATP + 14 NADH → 1 butanol | |
| ***Reductive Tricarboxylic Acid (rTCA) Cycle*** | | |
| $CO_2$ to metabolic intermediate | 4 $CO_2$ + 4 ATP + 8 NADH → 2 acetyl-CoA | Alissandratos *et al*. [100], Claassens *et al*. [101] |
| Intermediate to butanol | 2 acetyl-CoA + 4 NADH → 1 butanol | Schadeweg *et al*. [98] |
| *Overall $CO_2$ to butanol* | 4 $CO_2$ + 4 ATP + 12 NADH → 1 butanol | |
| ***3-hydroxypropionate/4-hydroxybutyrate (3HP-4HB) Pathway*** | | |
| $HCO_3^-$ to metabolic intermediate | 6 $HCO_3^-$ + 10 ATP + 10 NADH → 2 pyruvate | Berg *et al*. [99], Claassens *et al*. [98] |
| | 2 pyruvate → 2 acetyl-coA + 2 NADH + 2 $CO_2$ | Berg *et al*. [97], Schomburg *et al*. [94] |

| **Pathway** | **Reaction** | **Reference(s)** |
|---|---|---|
| Intermediate to butanol | 2 acetyl-CoA + 4 NADH → 1 butanol | Schadeweg *et al*. [98] |
| *Overall $HCO_3^-$ to butanol* | 6 $HCO_3^-$ + 10 ATP + 12 NADH → 1 butanol + 2 $CO_2$ | |
| ***3-hydroxypropionate (3HP) Cycle*** | | |
| $HCO_3^-$ to metabolic intermediate | 6 $HCO_3^-$ + 10 ATP + 12 NADH → 2 pyruvate | Zarzycki *et al*. [103], Herter *et al*. [104] |
| | 2 pyruvate → 2 acetyl-coA + 2 NADH + 2 $CO_2$ | Berg *et al*. [97], Schomburg *et al*. [94] |
| Intermediate to butanol | 2 acetyl-CoA + 4 NADH → 1 butanol | Schadeweg *et al*. [98] |
| *Overall $HCO_3^-$ to butanol* | 6 $HCO_3^-$ + 10 ATP + 14 NADH → 1 butanol + 2 $CO_2$ | |
| ***Formolase Pathway*** | | |
| $HCO_2^-$ to metabolic intermediate | 6 $HCO_2^-$ + 10 ATP + 4 NADH → 1 3-PG | Siegel *et al*. [65], Bar-Even *et al*. [66] |
| | 2 3-PG → 2 pyruvate + 2 ATP | Berg *et al*. [99] |
| | 2 pyruvate → 2 acetyl-coA + 2 NADH + 2 $CO_2$ | Berg *et al*. [97], Schomburg *et al*. [94] |
| | 2 acetyl-CoA + 4 NADH → 1 butanol | Schadeweg *et al*. [98] |
| *Overall $HCO_2^-$ to butanol* | 6 $HCO_2^-$ + 8 ATP + 6 NADH → 1 butanol + 2 $CO_2$ | |
| ***Dicarboxylate/4-hydroxybutyrate (4HB) Cycle*** | | |
| $CO_2$ to metabolic intermediate | 1 $CO_2$ + 1 $HCO_3^-$ + 3 ATP + 1 NADH + 6 Fd → 2 acetyl-CoA | Huber *et al*. [75] |
| Intermediate to butanol | 2 acetyl-CoA + 4 NADH → 1 butanol | Schadeweg *et al*. [98] |
| *Overall $CO_2$ to butanol* | 2 $CO_2$ + 2 $HCO_3^-$ + 6 ATP + 5 NADH + 12 Fd → 1 butanol | |

| **Thickness Range** | **Comment** | **Thickness (m)** | **References** |
|---|---|---|---|
| $h_1$ | *Geobacter sulfurreducens* biofilm | $4 \times 10^{-5}$ to $1.3 \times 10^{-4}$ | Malvankar *et al*. [59], Speers *et al*. [86] |
| $h_2$ | Polypyrrole conductive matrix for *Shewanella oneidensis* | $2 \times 10^{-4}$ to $6 \times 10^{-4}$ | Yu *et al*. [93] |
| $h_3$ | Cable bacteria biofilm and individual filaments | $1 \times 10^{-2}$ to $1.5 \times 10^{-2}$ | Pfeffer *et al*. [90], Meysman *et al*. [63] |

**Table S3.** Representative extracellular matrix thickness ranges considered in this article. Detailed discussion is included in **SI Text 9**. Reference numbers refer to main text references.

| **Resistivity Range** | **Comment** | **Resistivity (Ω cm)** | **References** |
|---|---|---|---|
| $\varrho_1$ | High conductivity polypyrrole | $1.1 \times 10^{-2}$ | Kang *et al*. [92] |
| $\varrho_2$ | *Thiofilum flexile* conductive filaments | $1.3 \times 10^{-2}$ | Meysman *et al*. [63] |
| $\varrho_3$ | Dried and fixed individual *Shewanella oneidensis* nanowires | 1 | El-Naggar *et al*. [62] |
| $\varrho_4$ | *Geobacter sulfurreducens* and *Shewanella oneidensis* biofilms | 200 to 8,952 | Malvankar *et al*. [59], Renslow *et al*. [60], Yates *et al*. [61] |
| $\varrho_5$ | Polypyrrole conductive matrix for *Shewanella oneidensis* | 312 | Yu *et al*. [93] |
| $\varrho_6$ | *E. coli* biofilm | $1.1 \times 10^{5}$ | Malvankar *et al*. [87] |
| $\varrho_7$ | HBr doped polyaniline | $2.17 \times 10^{6}$ | Hammo *et al*. [91] |
| $\varrho_8$ | Low conductivity polypyrrole | $1 \times 10^{9}$ | Kang *et al*. [92] |

**Table S4.** Representative extracellular matrix resistivity ranges considered in this article. Detailed discussion is included in **SI Text 9**. Reference numbers refer to main text references.

| Density Range | Comment | Densities (cells m$^{-3}$) | References |
|---|---|---|---|
| $n_1$ | Laboratory grown cultures of *E. coli* in exponential phase | $2 \times 10^{13}$ to $1 \times 10^{14}$ | BNIDs 103625 and 100985 |
| $n_2$ | Cyanobacteria grown to maximum density | $3.6 \times 10^{15}$ to $6.8 \times 10^{15}$ | Lippi *et al*. [81] |
| $n_3$ | Maximum density of laboratory grown *E. coli* | $5.1 \times 10^{15}$ to $1.39 \times 10^{17}$ | BNIDs 104940, 102231,100008, and 103904 |
| $n_4$ | *Ralstonia eutropha* grown under autotrophic conditions to maximum density | $1.0 \times 10^{17}$ to $2.0 \times 10^{17}$ | Tanaka *et al*. [42] |
| $n_5$ | Industrially grown microbes at maximum density | $1.5 \times 10^{18}$ | Abbas [82] |

**Table S5.** Representative cell density ranges considered in this article. Detailed discussion is included in **SI Text 5**. Reference numbers refer to main text references. BNID: BioNumbers ID.
